# Is FFT window length a neutral preprocessing choice in CNN-based dolphin whistle detection? A cross-domain sensitivity analysis

**DOI:** 10.64898/2026.05.01.721665

**Authors:** Rocco De Marco, Massimiliano Iurcev, Alessio Trebbi, Serena Lagorio

## Abstract

Convolutional neural networks (CNNs) operating on spectrogram images are the established method for automated cetacean whistle detection in passive acoustic monitoring (PAM). The FFT window length (*N*_*fft*_) used during spectrogram generation is routinely fixed without justification, yet it determines the time–frequency resolution and, consequently, how the resulting image is distorted when resized to a fixed CNN input dimension. This study presents a controlled sensitivity analysis of *N*_*fft*_ across five values (128, 256, 512, 1024, 2048) on binary *Tursiops truncatus* whistle detection, using 10-fold cross-validation on an in-domain dataset (Oltremare, 192 kHz) and cross-domain evaluation on an independent open-ocean benchmark (DCLDE 2022). All experiments were conducted within a formally defined, open-source pipeline (ai-pam-pipeline).

In-domain performance is uniformly high across all configurations. Cross-domain results diverge: *N*_*fft*_ = 256 significantly outperforms 512, 1024, and 2048 in macro F1, while maintaining a false discovery rate (FDR) of exactly zero across all 10 folds and all tested classification thresholds. *N*_*fft*_ = 128 achieves comparable recall but produces FDR > 0 in all 10 folds, exhibiting a transfer failure that is undetectable by in-domain validation. These results show that a preprocessing parameter routinely treated as an implementation detail has a large, systematic effect on cross-domain generalization. The study does not aim to propose *N*_*fft*_ = 256 as a universal optimum; rather, it tests the common assumption that this parameter is neutral and safe under a fully specified representation regime, and demonstrates that it is not.

## 1. Introduction

### 1.1 CNN-based detection in passive acoustic monitoring (PAM)

PAM is a widely-used technique for monitoring the presence and activity of marine mammals. Detecting the occurrence of these animals is an effective way to minimize the impact of human activities, particularly during potentially intrusive operations such as geophysical surveys involving multichannel seismic systems. Nowadays, marine mammal detection algorithms are increasingly automated and integrate artificial intelligence techniques to improve accuracy and efficiency. Therefore, understanding and optimizing the parameters that influence their performance is essential to enhance the reliability of PAM-based monitoring systems.

In recent years, convolutional neural networks (CNNs) operating on spectrogram representations have largely replaced traditional signal-processing techniques, such as template matching [1], active contours [2], and entropy-based methods [3] for automated cetacean vocalization detection in passive acoustic monitoring (PAM).

These performance gains are widely documented across detection [4,5], classification [6], segmentation [7], and transfer learning [8] tasks, and the approach now forms the backbone of most operational and research PAM workflows [9].

Less attention has been paid to the spectrogram itself. A survey of the CNN-based studies above reveals striking heterogeneity in input representations — image sizes ranging from 40 × 40 to 1000 × 1000 pixels, single-channel or three-channel formats, and STFT windows lengths ranging from a few milliseconds to over 100 ms. In every case, these parameters were fixed *a priori* and held constant throughout the study. Li et al. [6], in one of the few explicit acknowledgments of this issue, noted that their chosen parameters were “not necessarily the optimal spectrogram specification,” yet did not pursue the question further. Stowell [9] identified preprocessing standardization as an open challenge for computational bioacoustics more broadly.

This inattention contrasts with the wider audio machine-learning literature, where the choice of time–frequency representation is itself treated as a key subject of study. In polyphonic music transcription, for example, Cheuk et al. [10] fixed the network architecture and varied only the input representation — STFT window length, number of frequency bins, and frame length — obtaining an 8.3% change in transcription accuracy from this choice alone, and reporting the counter-intuitive result that the highest spectral resolution does not always yield the best performance. Comparable representation-sensitivity studies exist for environmental sound and acoustic scene classification.

In bioacoustic signal processing, selecting *N*_fft_ has traditionally been framed as an unavoidable trade-off between temporal and spectral resolution [11]. However, while human annotators visually adapt to varying time–frequency trade-offs when inspecting spectrograms, deep learning models learn features directly bounded by the fixed grid geometry imposed during feature extraction [12]. Furthermore, when time–frequency matrices are resized to match fixed CNN input dimensions (e.g., 224 × 224 or 256 × 256), different *N*_fft_ values introduce non-uniform spatial interpolation artifacts [13]. This effectively alters the aspect ratio and slope of vocalization contours, forcing the network to learn representation-dependent features rather than invariant acoustic patterns.

While recent advances in general audio ML have proposed learnable time–frequency front-ends to bypass manual STFT parameter selection [14, 15], standard STFT spectrograms remain the operational standard in PAM due to their computational efficiency, interpretability, and seamless integration into established bioacoustic software.

In cetacean PAM, however, the analogous question remains critically unaddressed: specifically, how the FFT window length (*N*_*fft*_) affects CNN detection performance, and whether that effect scales to novel recording environments.

### 1.2 The FFT window length as an unexamined hyperparameter

The FFT window length (*N*_*fft*_) controls the time–frequency trade-off of the STFT. At a given sampling rate, it determines the number of frequency bins that represent a vocalization and, consequently, how that representation is distorted when resized to a fixed-dimension CNN input image. A typical bottlenose dolphin whistle spanning 5–20 kHz occupies frequency bins 7-27 at *N*_*fft*_ = 256 (192 kHz, 750 Hz/bin), but spans bins 53-213 at *N*_*fft*_ = 2048 (94 Hz/bin). When both spectrograms are resized to 224 × 224 pixels, the geometric transformation applied to the whistle trace is qualitatively different: the first is upsampled into a thick, high-contrast band; the second is compressed into a thin contour embedded in a richer noise structure. Despite this predictable interaction, no published study treats *N*_*fft*_ as an independent experimental variable with controlled cross-domain evaluation in a cetacean PAM context.

### 1.3 Cross-domain evaluation and the limits of in-domain validation

The distinction between in-domain and cross-domain performance is central to this study. A detector validated on held-out data from the same recording session provides an optimistic upper bound; operational deployment invariably involves different recording equipment, acoustic environments, noise profiles, and — for highly mobile marine species — different populations and individual vocal repertoires. This distribution shift between training and deployment has been identified as one of the defining challenges of bioacoustic machine learning [17], where models that appear well-calibrated in-domain can degrade sharply on out-of-distribution data. A parameter whose effect is negligible in-domain but decisive cross-domain would, by construction, escape detection under the validation protocols most commonly reported in the PAM literature. The FFT window length (*N*_*fft*_), as this study will show, is exactly such a parameter.

### 1.4 Objectives

This study presents a controlled sensitivity analysis of *N*_*fft*_ across five values (128, 256, 512, 1024, 2048) on binary *T. truncatus* whistle detection. All experiments were conducted within the six-stage workflow formalized in [18] and implemented in the open-source toolkit ai-pam-pipeline [19], which ensures exact reproducibility through a single YAML configuration file. Cross-domain evaluation was performed on an independent open-ocean dataset (DCLDE 2022 [20]). A classification-threshold sweep across five operating points examines the robustness of the finding; a supplementary probe under a different image geometry, reported in Supplementary Material, tests whether the ranking generalizes beyond the primary input dimensions. The study does not propose a universal optimum; it tests whether the neutrality assumption is safe, under one fully specified representation regime.

## 2. Methods

### 2.1 Datasets

#### Oltremare (in-domain)

The training corpus consists of continuous recordings of seven *Tursiops truncatus* at the Oltremare facility (Riccione, Italy), acquired in November 2021 at 192 kHz; the recording setup and annotation procedure are described in Screpanti et al. [21] and in the framework article [18]. From these recordings, 3,451 whistle and 233 multiple-whistle annotations form the positive class (C1); the negative class (C0) comprises background noise, echolocation click trains, burst-pulse sounds, and feeding buzzes — the same realistic composition adopted in [18], which forces the detector to reject co-occurring non-target vocalizations rather than only silence. The annotated dataset is publicly available [22] and has been used in [16,18] for different experimental purposes.

#### DCLDE 2022 (cross-domain)

Cross-domain evaluation relied on a *Tursiops truncatus* encounter from the HICEAS 2017 towed-array survey in the central Pacific [20]. The encounter (~15 minutes, detection 46) was selected because it is the only publicly available open-ocean *T. truncatus* recording with temporal annotations independent of the Oltremare corpus. A balanced test set of 500 whistle and 500 non-whistle spectrograms was drawn from the annotated pool (randomly selected with a given seed = 42); no DCLDE material entered the training pipeline. The same encounter appears in [16] for a filtering comparison at fixed *N*_*fft*_ = 512 and in [18] for a single-configuration cross-domain evaluation; the present study extends its use to the full five-value *N*_*fft*_ sweep.

### 2.2 Pipeline overview

Experiments followed the six-stage workflow formalized in [18] and implemented in the open source toolkit ai-pam-pipeline [19]. The pipeline progresses from audio acquisition and PAM annotation (Stage 1) through segmentation and windowing (Stage 2), spectrogram generation (Stage 3), optional image preprocessing (Stage 4), CNN training with k-fold cross-validation (Stage 5), to model evaluation (Stage 6). In the present study, no image-domain filtering was applied (Stage 4 was limited to per-image intensity normalization).

To clarify the signal-to-image transformation process: an audio segment of 0.8 s is first converted into a time-frequency representation via STFT. The native dimensions of this resulting matrix depend strictly on the chosen *N*_*fft*_ and the STFT window overlap (fixed at 0.5). Following the STFT, the matrix is cropped along the frequency axis to retain only the 3–25 kHz band. As the CNNs require a fixed-dimension input, this variable-sized native matrix is then geometrically resized to the target image resolution (224 × 224 pixels for the primary arm, or 300 × 150 for the geometry probe) using bilinear interpolation. Finally, in Stage 4, per-image intensity normalization (min-max scaling to a [0, 1] range) is applied before feeding the spectrogram to the network.

Detailed information about the pipeline is provided in the Extended Data accompanying the framework article [23].

### 2.3 CNN architecture

The detector is the lightweight three-block CNN introduced by Scaradozzi et al. [24] for edge deployment and subsequently used in [16,18]: filter progression 32, 64, 128, each block with ReLU and 2 × 2 pooling, followed by a 128-unit dense layer and a sigmoid output. The input is a single-channel grayscale spectrogram (224 × 224 in the primary arm; 300 × 150 in the geometry probe). Training strictly followed the protocol documented in [18]: Adam (lr = 10_−4_), batch size of 32, early stopping with a patience of 10 epochs (including a 5-epoch warmup) and best-weight restoration. The architecture was deliberately kept fixed across all 150 runs; the objective of this study is to isolate the effect of the preprocessing parameter, rather than to optimize the classifier.

### 2.4 Experimental design

The primary independent variable was *N*_*fft*_ ∈ {128, 256, 512, 1024, 2048}. One additional experimental dimension was included as a sensitivity check:

- **Overlap condition:** as_overlap ∈ {0.5, 0.0}. The primary arm used 0.5 (the pipeline’s scroll-augmentation mode); the check at 0.0 rules out augmentation-from-overlap as a confound for the *N*_*fft*_ ranking.

A supplementary probe replicating the full *N*_*fft*_ sweep under a 300 × 150 image geometry, shown by [16] to outperform 224 × 224 at fixed *N*_*fft*_ = 512, was also run; full results are reported in Supplementary Material (SM-3) and discussed briefly in Section 4.4.

All other parameters were held constant: analysis window length = 0.8 s, STFT overlap = 0.5, frequency crop = 3–25 kHz, kernel = 3 × 3, 10-fold stratified cross-validation, seed = 42, classification threshold θ = 0.5. The primary arm (as_overlap = 0.5) used 4,700 samples per class; the overlap-check arm (as_overlap = 0.0) used 3,000, determined by the binding class count at zero overlap. A total of 15 training configurations (5 *N*_*fft*_ × 2 overlap conditions + 5 *N*_*fft*_ × geometry probe) yielded 150 independently trained fold models; primary-arm results are reported in the main text, with the full grid in Supplementary Material.

Table 1 summarizes the experimental grid. Complete YAML configurations for all 15 training runs and 10 crosstest generation runs are provided as Supplementary Material; the YAML schema is fully documented in *ai-pam-pipeline* article [16] and in the Supplementary Material.

**Table 1.** Experimental grid. Cells above the line are variables; cells below the line are constants shared across all 150 fold models.

| Parameter | Values |
| --- | --- |
| $N_{fft}$ (independent variable) | 128, 256, 512, 1024, 2048 |
| Window overlap (sensitivity check) | 0.5 (primary), 0.0 (check) |
| Image geometry (probe) | $224 \times 224$ (primary), $300 \times 150$ (probe, ov = 0.5 only) |
| Classification threshold $\theta$ (sweep) | 0.1, 0.3, 0.5, 0.7, 0.9 |
| Analysis window | 0.8 s |
| STFT overlap | 0.5 |
| Frequency crop | 3–25 kHz |
| CNN Architecture | baseline_cnn.py ($3 \times 3$ kernel) [24] |
| k-fold CV | 10 folds, seed = 42 |
| Samples/class | 4,700 (ov = 0.5) / 3,000 (ov = 0.0) |

### 2.5 Statistical methodology

Statistical significance was assessed using the paired Wilcoxon signed-rank test [25], applied to the macro F1 and recall values obtained from each of the 10 folds. In the primary analysis, four pairwise comparisons were conducted (256 vs. each of {128, 512, 1024, 2048}), with Bonferroni correction yielding α_adj_ = 0.05/4 = 0.0125. A full 10-pairwise comparison matrix (α_adj_ = 0.05/10 = 0.005) is reported in the Supplementary Material. The rank-biserial correlation (*r*) was computed as effect size, where |*r*| > 0.5 is interpreted as a large effect. These tests were performed independently for in-domain and cross-domain results.

### 2.6 Threshold analysis

For each of the five *N*_*fft*_ configurations, the ten fold models were evaluated on the DCLDE cross-domain dataset across the threshold range θ ∈ {0.10, 0.30, 0.50, 0.70, 0.90}. At each threshold value, recall, macro F1, and FDR were computed on the balanced test set (500 samples per class). The threshold sweep was also replicated at as_overlap = 0.0 to verify cross-arm consistency; results from the 300 × 150 geometry probe are reported in Supplementary Material (SM-4). A threshold-invariance index ΔF1 = F1_max_ − F1_min_ over the θ range is reported for each *N*_*fft*_ as a summary measure of operational stability. Self-check: the θ = 0.5 values were verified against the existing cross test results across all 150 fold models (maximum discrepancy = 0.00).

### 2.7 Error reporting convention

All classifier outputs flagged as false positives were retained as such throughout the analysis, without post-hoc reclassification. An isolated, uncorroborated acoustic trace — even where faintly whistle-compatible — does not meet the labeling standard applied elsewhere in this study, particularly given the operational cost of a false positive in a deterrence context; this conservative convention is applied uniformly across all *N*_*fft*_ configurations and datasets.

## 3. Results

### 3.1 In-domain performance

Table 2 and Figure 1 report the in-domain validation performance for the primary arm (as_overlap = 0.5, 224 × 224). The five *N*_*fft*_ configurations are virtually indistinguishable: macro F1 spans a 0.020 range (0.955–0.975) and no pairwise Wilcoxon comparison reaches significance (all *p* > 0.05). On this evidence alone, a practitioner would reasonably conclude that *N*_*fft*_ does not matter, a conclusion that the cross-domain data will overturn.

**Table 2.** In-domain cross-validation performance (Oltremare, θ = 0.5, primary arm): mean ± SD across 10 folds.

| $N_{fft}$ | Accuracy | Macro F1 | FDR |
| --- | --- | --- | --- |
| 128 | $0.968 \pm 0.006$ | $0.968 \pm 0.006$ | $0.022 \pm 0.008$ |
| **256** | **$0.975 \pm 0.005$** | **$0.975 \pm 0.005$** | **$0.015 \pm 0.007$** |
| 512 | $0.969 \pm 0.008$ | $0.969 \pm 0.008$ | $0.022 \pm 0.014$ |
| 1024 | $0.961 \pm 0.010$ | $0.961 \pm 0.010$ | $0.030 \pm 0.016$ |
| 2048 | $0.955 \pm 0.021$ | $0.955 \pm 0.021$ | $0.033 \pm 0.013$ |

**Figure 1.**
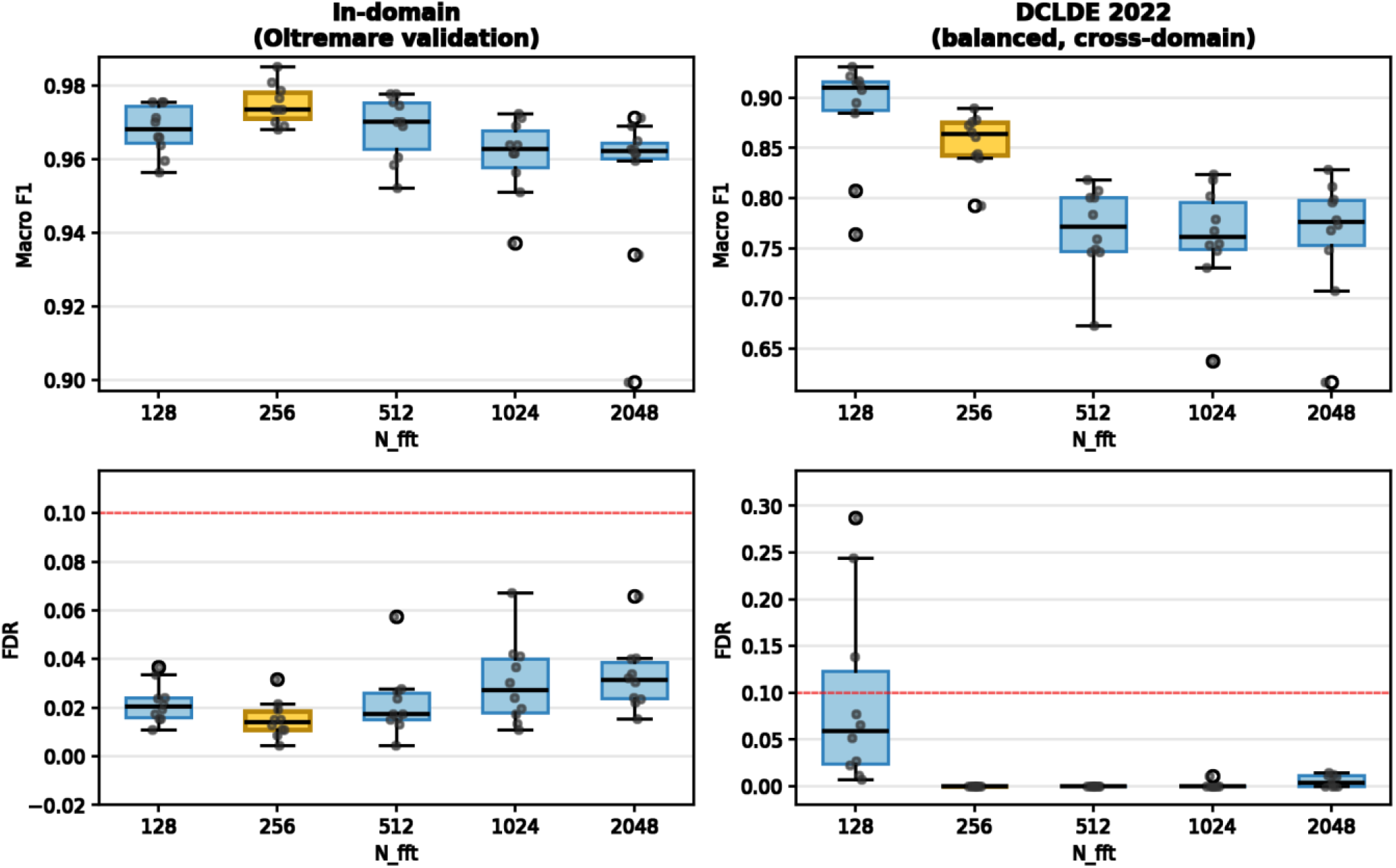
Per-fold performance comparison, model 224×224, spectrogram overlap 50%, 10 folds. Left: in domain F1 and Recall results, on right Cross-domain (DCLDE 2022) results.

### 3.2 Cross-domain performance

The cross-domain evaluation on the DCLDE dataset reveals the effect that in-domain metrics conceal. Table 3 and Figure 1 report the results for the primary arm.

**Table 3.** Cross-domain performance (DCLDE 2022, θ = 0.5, primary arm): mean ± SD across 10 folds. Wilcoxon *p*-values are Bonferroni-corrected (α_adj_ = 0.0125).

| $N_{fft}$ | Recall | Macro F1 | FDR | vs. 256: F1 $p$ | vs. 256: $r$ |
| --- | --- | --- | --- | --- | --- |
| 128 | $0.886 \pm 0.052$ | $0.886 \pm 0.052$ | $0.093 \pm 0.094$ | 0.275 (ns) | — |
| **256** | **$0.860 \pm 0.025$** | **$0.856 \pm 0.026$** | **$0.000 \pm 0.000$** | — | — |
| 512 | $0.781 \pm 0.035$ | $0.769 \pm 0.041$ | $0.000 \pm 0.000$ | 0.002** | 1.000 |
| 1024 | $0.775 \pm 0.042$ | $0.762 \pm 0.051$ | $0.001 \pm 0.003$ | 0.002** | 1.000 |
| 2048 | $0.776 \pm 0.047$ | $0.763 \pm 0.058$ | $0.006 \pm 0.006$ | 0.002** | 1.000 |

$N_{fft} = 256$ significantly outperforms 512, 1024, and 2048 on both macro F1 and recall, with maximum effect size ( $r = 1.0$ ): every single fold of 256 exceeds every single fold of the three longer-window configurations, without exception.
$N_{fft} = 128$ achieves numerically higher recall (0.886 vs. 0.860) but with double the inter-fold standard deviation and, critically, a FDR of 0.093, driven by false-positive bursts in individual folds (e.g., fold 6: 181 FP out of 500 noise samples). The Wilcoxon test on FDR confirms a significant difference between 256 and 128 ( $p = 0.002$ , $r = -1.0$ ).
$N_{fft} = 128$ produces $FDR > 0$ in 10 out of 10 folds, while $N_{fft} = 256$ produces $FDR = 0.000$ in 10 out of 10 folds.
The full 10-pairwise Wilcoxon matrix (Supplementary Table SM-1) confirms a two-tier structure: {128, 256} significantly outperform {512, 1024, 2048} on recall and F1, while the three longer-window configurations are mutually indistinguishable ( $p > 0.8$ for all pairwise comparisons). Within the top tier, 256 separates from 128 on specificity alone.

### 3.3 Overlap sensitivity check

The *N*_*fft*_ ranking was preserved when the scroll-augmentation overlap was removed (as_overlap = 0.0): *N*_*fft*_ = 256 achieved F1 = 0.819, followed by 512 (0.757), 2048 (0.730), and 1024 (0.717). *N*_*fft*_ = 128 achieved F1 = 0.897 with FDR = 0.056 — lower than in the primary arm but still non-zero. The consistency of the ranking across overlap conditions rules out augmentation from overlapping windows as a confound. The full table is reported in Supplementary Material (SM-2).

### 3.4 Threshold sensitivity

Table 4 and Figure 2 report the threshold sweep results for the primary arm. Three patterns emerge from the 5 × 5 matrix.

**Table 4.** Threshold sensitivity — cross-domain macro F1 and FDR as a function of θ and *N*_*fft*_ (primary arm, mean ± SD over 10 folds).

Macro F1:
| $N_{fft}$ $\theta$ | 0.1 | 0.3 | 0.5 | 0.7 | 0.9 |
| --- | --- | --- | --- | --- | --- |
| 128 | 0.681 ± 0.155 | 0.840 ± 0.103 | 0.886 ± 0.052 | 0.897 ± 0.027 | 0.889 ± 0.022 |
| **256** | **$0.899 \pm 0.021$** | **$0.872 \pm 0.025$** | **$0.856 \pm 0.026$** | **$0.838 \pm 0.030$** | **$0.810 \pm 0.031$** |
| 512 | $0.847 \pm 0.029$ | $0.801 \pm 0.037$ | $0.769 \pm 0.041$ | $0.738 \pm 0.047$ | $0.691 \pm 0.058$ |
| 1024 | $0.849 \pm 0.045$ | $0.804 \pm 0.045$ | $0.762 \pm 0.051$ | $0.719 \pm 0.052$ | $0.656 \pm 0.066$ |
| 2048 | $0.850 \pm 0.028$ | $0.804 \pm 0.045$ | $0.763 \pm 0.058$ | $0.723 \pm 0.070$ | $0.654 \pm 0.088$ |

FDR:
| $N_{fft}$ $\theta$ | 0.1 | 0.3 | 0.5 | 0.7 | 0.9 |
| --- | --- | --- | --- | --- | --- |
| 128 | $0.336 \pm 0.106$ | $0.168 \pm 0.126$ | $0.093 \pm 0.094$ | $0.045 \pm 0.062$ | $0.011 \pm 0.022$ |
| **256** | **$0.001 \pm 0.002$** | **$0.001 \pm 0.001$** | **$0.000 \pm 0.000$** | **$0.000 \pm 0.000$** | **$0.000 \pm 0.000$** |
| 512 | $0.007 \pm 0.014$ | $0.000 \pm 0.001$ | $0.000 \pm 0.000$ | $0.000 \pm 0.000$ | $0.000 \pm 0.000$ |
| 1024 | $0.032 \pm 0.071$ | $0.006 \pm 0.018$ | $0.001 \pm 0.003$ | $0.000 \pm 0.000$ | $0.000 \pm 0.000$ |
| 2048 | $0.054 \pm 0.041$ | $0.016 \pm 0.014$ | $0.006 \pm 0.006$ | $0.001 \pm 0.003$ | $0.001 \pm 0.003$ |

First, the $N_{fft}$ ranking is threshold-invariant: $N_{fft} = 256$ achieves the highest F1 at every threshold except $\theta = 0.7$ and $\theta = 0.9$ , where $N_{fft} = 128$ matches or slightly exceeds it — but only by sacrificing specificity (FDR = 0.045 and 0.011, respectively, vs. 0.000 for 256).
$N_{fft} = 256$ is the most threshold-stable configuration: its F1 range across the full $\theta$ sweep ( $\Delta F1 = 0.089$ ) is less than half that of $N_{fft} = 128$ ( $\Delta F1 = 0.216$ ) and substantially smaller than 512 (0.156), 1024 (0.193), or 2048 (0.196).
$N_{fft} = 128$ escalates sharply at low thresholds, reaching 0.336 at $\theta = 0.1$ — a rate at which one in three positive detections is a false alarm.
$N_{fft} = 256$ , by contrast, maintains $FDR \leq 0.001$ across the entire threshold range.
This pattern replicates at $as\_overlap = 0.0$ , where the $\Delta F1$ for 256 is 0.104 (vs. 0.184 for 128). The $300 \times 150$ geometry probe confirms the same ranking, with full cross-arm threshold matrices provided in Supplementary Material (SM-3, SM-4).

**Figure 2.**
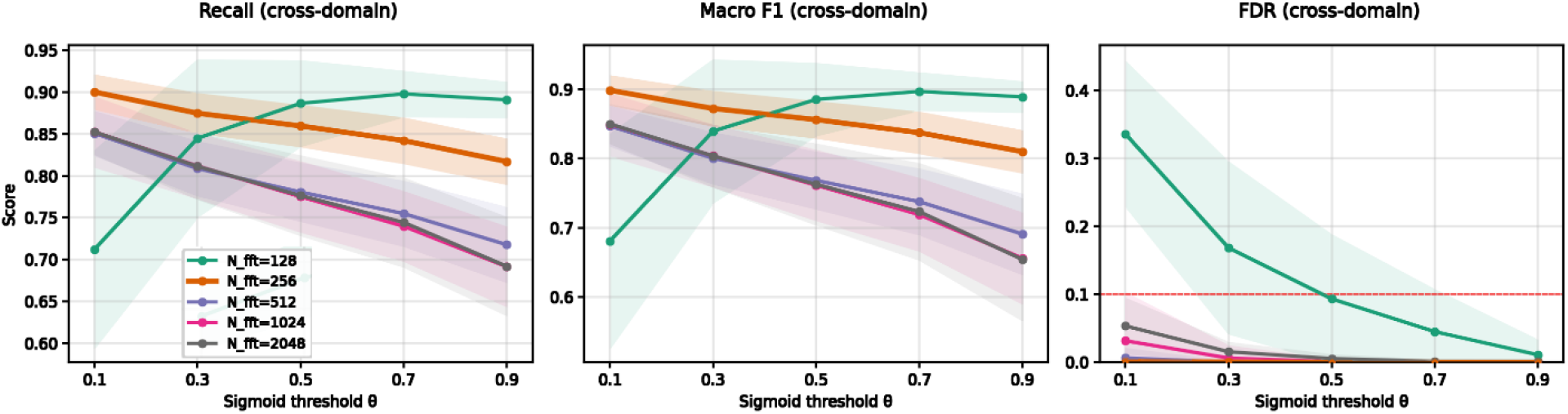
Threshold sensitivity analysis versus DCLDE 2022 Cross-domain dataset, model 224×224, spectrograms overlap 0.5. Recall, Macro F1 and FDR have been compared across the different *N*_*fft*_

First, the *N*_*fft*_ ranking is threshold-invariant: *N*_*fft*_ = 256 achieves the highest F1 at every threshold except θ = 0.7 and θ = 0.9, where *N*_*fft*_ = 128 matches or slightly exceeds it — but only by sacrificing specificity (FDR = 0.045 and 0.011, respectively, vs. 0.000 for 256).

*N*_*fft*_ = 256 is the most threshold-stable configuration: its F1 range across the full θ sweep (ΔF1 = 0.089) is less than half that of *N*_*fft*_ = 128 (ΔF1 = 0.216) and substantially smaller than 512 (0.156), 1024 (0.193), or 2048 (0.196).

*N*_*fft*_ = 128 escalates sharply at low thresholds, reaching 0.336 at θ = 0.1 — a rate at which one in three positive detections is a false alarm.

*N*_*fft*_ = 256, by contrast, maintains FDR ≤ 0.001 across the entire threshold range.

This pattern replicates at as_overlap = 0.0, where the ΔF1 for 256 is 0.104 (vs. 0.184 for 128). The 300 × 150 geometry probe confirms the same ranking, with full cross-arm threshold matrices provided in Supplementary Material (SM-3, SM-4).

## 4. Discussion

### 4.1 *N*_*fft*_ = 256 as the Pareto-optimal configuration

The central finding is that *N*_*fft*_ = 256 occupies a unique position in the parameter space: it is the only configuration that combines high cross-domain recall with structurally zero false discovery rate across all thresholds and all experimental arms. The relationship between *N*_*fft*_ and cross-domain performance is not monotonic. Shorter windows do not uniformly improve generalization: *N*_*fft*_ = 128 produces numerically higher recall than 256 at θ = 0.5 (0.886 vs. 0.860), but at the cost of a false-positive rate that no operational deployment could tolerate, particularly in automated deterrence systems where each positive detection triggers an intervention.

Limiting the activation of deterrence mechanisms—typically involving the emission of disruptive acoustic signals—is a primary objective for preserving marine ecosystems, even at the expense of occasional missed detections. *N*_*fft*_ = 256, by contrast, exhibits FDR = 0.000 across all 50 fold × threshold combinations (10 folds × 5 θ values) in the primary arm, a level of specificity that is structurally absent from any other configuration.

At the other end of the spectrum, the three longer-window configurations (512, 1024, 2048) are mutually indistinguishable in cross-domain performance (Wilcoxon *p* > 0.8 for all pairwise comparisons) but consistently and significantly inferior to 256 on both recall and F1. The practical recommendation is therefore specific: at 192 kHz, for tonal cetacean vocalizations analyzed with this class of lightweight CNN on fixed-dimension spectrogram images, *N*_*fft*_ = 256 is a reasonable starting point. If the sampling rate, the target vocalization, or the image dimensions differ, the parameter should be tested explicitly rather than assumed.

### 4.2 Physical mechanism

The performance differences across *N*_*fft*_ values can be understood through the interaction between spectral resolution and image resize (Figure 3). At 192 kS/s, a *T. truncatus* whistle (2– 10 kHz) occupies the following number of frequency bins depending on *N*_*fft*_: 1–6 bins at *N*_*fft*_ = 128 (1500 Hz/bin), 3–13 at 256 (750 Hz/bin), 5–27 at 512 (375 Hz/bin), 10–53 at 1024 (188 Hz/bin), and 21–106 at 2048 (94 Hz/bin). When the raw spectrogram is resampled to a fixed 224 × 224 pixel grid, the geometric transformation applied to the whistle trace differs qualitatively across these conditions.

**Figure 3.**
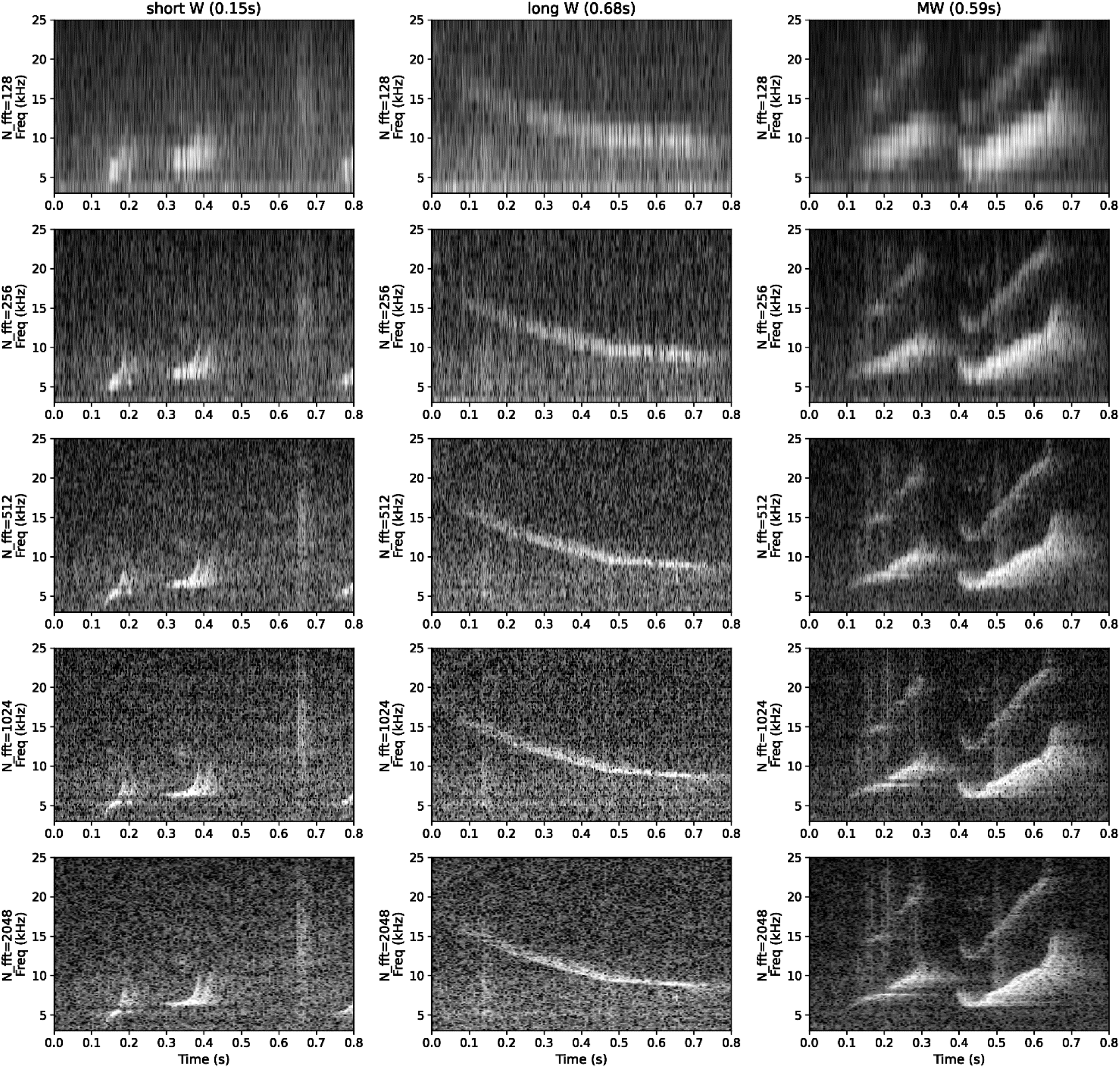
Spectrogram visualization of three different whistles (short, long and multiple) for different *N*_*fft*_ settings.

At *N*_*fft*_ = 256, the bilinear upsampling acts as an implicit spatial amplifier: the signal energy, concentrated in a small number of bins, is spread over a wide bright band that occupies a large fraction of the image height. A consequence is a significant increase in local contrast, the intensity gradient at the trace boundary relative to the surrounding noise floor is steep. At *N*_*fft*_ = 1024 or 2048, the same acoustic energy is distributed over many more frequency bins; each bin receives fewer pixels after upsampling, reducing per-pixel intensity and local contrast. The result is a thinner whistle contour embedded in a richer noise structure that carries more recording-specific texture and is therefore less transferable across domains. Additionally, a shorter FFT window produces more temporal frames for a fixed analysis window (approximately 1,200 frames at *N*_*fft*_ = 256 vs. 150 at *N*_*fft*_ = 2048), providing the CNN with finer temporal sampling of the FM trajectory.

These three concurrent effects — wider trace, higher local contrast, finer temporal sampling — collectively favor shorter FFT windows for cross-domain generalization. However, *N*_*fft*_ = 128 pushes the representation past the point of diminishing returns: with only 1–6 bins per whistle, the FM contour lacks sufficient frequency detail for the CNN to learn a stable, generalizable feature. The result is a model that memorizes recording-specific intensity patterns — patterns shared by the training and validation sets (both drawn from the same Oltremare session) but absent from the DCLDE recording.

### 4.3 In-domain evaluation is blind to the *N*_*fft*_ effect

The in-domain results (Table 2) present a uniform picture: five configurations, 0.020 spread in macro F1, no significant pairwise differences. A researcher who validates exclusively on held-out folds from the same recording session — as is common practice in the PAM literature — would have no reason to suspect that *N*_*fft*_ matters. Yet the cross-domain data show an F1 range of 0.124 (0.762–0.886) across the same five configurations. The discrepancy arises because training and validation folds share the recording-level acoustic structure of Oltremare; any feature that the CNN extracts from that structure will score well in-domain regardless of whether it transfers to a different environment.

The case of *N*_*fft*_ = 128 illustrates the point with particular clarity. Fold 6 of *N*_*fft*_ = 128 achieves the lowest validation loss in the entire 150-model battery (0.071) and the smallest generalization gap between training and validation loss (0.087), yet it produces the worst cross-domain performance of any fold in any configuration (F1 = 0.764, FDR = 0.287). By contrast, fold 6 of *N*_*fft*_ = 256 exhibits a larger generalization gap (0.152) and a steeper validation-loss trend over its final 10 epochs (+0.087), indicators that conventional diagnostics would flag as concerning, yet it is entirely stable cross-domain (F1 = 0.878, FDR = 0.000). The in-domain training dynamics do not predict cross-domain FDR. The failure of *N*_*fft*_ = 128 is not classical overfitting; it is a transfer failure tied to spectral resolution, in which the CNN learns features specific to the recording-level noise structure shared between training and validation folds (both drawn from the same Oltremare session) that do not transfer to the acoustically distinct DCLDE environment. Full training histories and generalization-gap metrics for these folds are provided in Supplementary Material (SM-7).

This finding carries a methodological implication beyond the scope of *N*_*fft*_ selection: cross-domain evaluation on independently recorded data should be a standard component of PAM detector validation, not an optional supplement. In-domain k-fold cross-validation, even when carefully implemented, cannot diagnose representation-level failures that manifest only under domain shift.

### 4.4 Directions for further parametrization

The geometry probe provides direct evidence for a further axis of the mechanism. Scaradozzi et al. [16] reported that 300 × 150 spectrograms outperform 224 × 224 at fixed *N*_*fft*_ = 512, attributing the advantage to improved temporal context. The present data confirm the observation, 300 × 150 outperforms 224 × 224 at every tested *N*_*fft*_, but reveal that the magnitude of the gap is *N*_*fft*_-dependent: ΔF1 = +0.012 at *N*_*fft*_ = 128, +0.043 at 256, +0.087 at 512, +0.063 at 1024, and +0.050 at 2048 (Supplementary Table SM-3). The gap is largest precisely where the square resize introduces the most distortion, at intermediate *N*_*fft*_ values where the native spectrogram aspect ratio differs most from a square, and smallest at 128, where so few bins exist that even a less distortive resize cannot compensate for insufficient spectral resolution. This interaction is inconsistent with a generic “temporal context” explanation and instead points to the resize distortion of the native bin structure as the primary driver. Under the 300 × 150 geometry, *N*_*fft*_ = 256 achieves F1 = 0.899 with FDR = 0.000, the highest reliable operating point in the battery, while *N*_*fft*_ = 128 reaches a numerically comparable F1 (0.898) but with FDR = 0.082, preserving the specificity gap observed at 224 × 224.

The interaction between *N*_*fft*_ and image geometry can be summarized by two derived quantities. Given sampling rate *f*_*s*_, frequency crop [*f*_*min*_, *f*_*max*_], and image dimensions *H* × *W*, the native number of frequency bins within the crop is *N*_*bins*_ = (*f*_*max*_ − *f*_*min*_)*N*_*fft*_/*f*_*s*_, and the native number of STFT frames spanning a segment of duration *T*_*seg*_ is *N*_*frames*_ = *T*_*seg*_*f*_*s*_/[*N*_*fft*_(1 − *ov*_*stft*_)], where *ov*_*stft*_ is the STFT frame overlap (fixed at 0.5 throughout this study; distinct from the segmentation-window overlap varied in Section 3.3). The image-space representation density along each axis is then

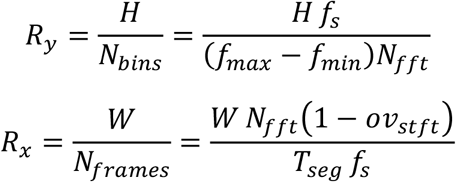

pixels of rendered image per native frequency bin (*R*_y_) or STFT frame (*R*_x_). The two move in opposite directions as *N*_*fft*_ grows: *R*_y_ falls (frequency-axis compression) while *R*_x_ rises (time-axis stretching), so a geometric account of the mechanism must consider both jointly rather than in isolation. Both are offered as descriptive devices, not independently validated predictors (full derivation, including the final-layer analogues *R*_y,last_ and *R*_x,last_, in Supplementary Material SM-10).

A tempting simplification — that the optimal *N*_*fft*_ is the one bringing the network’s final-layer analogue *R*_y,last_ closest to unity — does not hold consistently: for the primary 224 × 224 geometry (*H*_last_ = 26) it correctly identifies *N*_*fft*_ = 256, but applied to the 300 × 150 geometry probe (*H*_last_ = 17) it would predict *N*_*fft*_ = 128, whereas the empirical data above show *N*_*fft*_ = 256 remains superior there too. This is a useful negative result: the interaction between spectral resolution and network architecture is not reducible to a single geometric ratio, even one computed at the network’s own output resolution — reinforcing, rather than undermining, the caution below against treating *N*_*fft*_ as portable across representation regimes.

*R*_x_ and *R*_y_ describe only the geometric side of the interaction. Network architecture — depth and layer width in particular — is a further axis: spectral resolution and network capacity jointly determine what information reaches and is exploitable by the classifier, so the ranking reported here is not guaranteed to transfer to networks of different depth or width. *N*_*fft*_ should therefore be regarded as one axis of a larger joint design space — sampling rate, image geometry, and network capacity together — rather than a hyperparameter that can be tuned once and reused across architectures. This does not diminish its relevance: on the evidence presented here, *N*_*fft*_ remains one of the most consequential of these design choices, and one of the least examined.

### 4.5 Scope statement

CNN-based classification of cetacean vocalizations is an intrinsically multi-parametric problem — segmentation windowing, spectrogram resolution, image geometry, filtering, architecture depth, and sampling rate all interact, which is precisely why ai-pam-pipeline was built to make each choice explicit and reproducible rather than implicit. Expanding this study to jointly explore that full parameter space would risk an unbounded search with diminishing interpretability. This study isolates one parameter — *N*_*fft*_ — that has received no dedicated attention despite being fixed by convention in every reviewed study and shows that the convention is not neutral. The intention is not to resolve the parametrization of cetacean whistle detection, but to demonstrate that it requires resolving, and to provide a reproducible template for doing so.

### 4.6 Threshold invariance, diagnostic value, and practical recommendation

The FDR of *N*_*fft*_ = 256 remains at or below 0.001 across the entire θ range (0.1–0.9) in all three experimental arms. This property is operationally valuable in PAM deployments where the detector output drives automated responses — acoustic deterrence devices, mitigation protocols, or real-time survey adjustments — because it means the threshold can be tuned freely to balance recall against missed-detection cost without introducing false alarms.

*N*_*fft*_ = 128, by contrast, reaches FDR = 0.336 at θ = 0.1, a rate at which one in three positive detections would trigger an unnecessary response. Even at the balanced operating point (θ = 0.5), its FDR of 0.093 exceeds what most operational systems would accept.

The threshold-invariance index ΔF1 provides a compact summary of detector operational stability.

For *N*_*fft*_ = 256, ΔF1 ranges from 0.089 to 0.104 across the three experimental arms — the smallest of any configuration.

For *N*_*fft*_ = 128, the range is 0.184–0.308, two to three times larger, indicating that its performance is strongly threshold-dependent. This metric may serve as a useful complement to point-estimate performance measures (F1 at a single θ) when comparing detectors intended for deployment under varying operational constraints.

Beyond its role in operational tuning, where a stricter or more permissive θ trades recall against false-discovery rate depending on deployment context, the shape of the θ-sensitivity curve carries diagnostic value. A configuration whose FDR falls substantially as θ is raised indicates that the underlying score distributions for whistle and non-whistle windows are separable, and that the default θ = 0.5 was simply a miscalibrated operating point for that configuration; a configuration whose FDR remains elevated across the entire θ range indicates a more fundamental failure of class separability that threshold adjustment alone cannot repair. *N*_*fft*_ = 128 shows substantial recovery at higher θ (FDR falling from 0.336 at θ = 0.1 to 0.011 at θ = 0.9) yet never approaches the near-zero FDR that *N*_*fft*_ = 256 sustains without any threshold adjustment, consistent with a partially recoverable, calibration-level effect for 128 layered on top of a more fundamental representational limitation at that window length. Threshold sensitivity analysis is therefore recommended not only for setting a deployment operating point, but as a diagnostic of whether a given *N*_*fft*_ configuration has converged to a genuinely discriminative representation.

For 192-kHz bottlenose dolphin whistle detection using 0.8-s windows and a 224 × 224 spectrogram input with this class of lightweight CNN, *N*_*fft*_ = 256 gives the best tested recall–FDR trade-off and is a reasonable default to test first. The broader conclusion is that *N*_*fft*_ should be included in the parameter space explicitly evaluated when developing a new detector, rather than fixed by convention.

### 4.7 Limitations

The experimental evidence presented here is subject to several constraints, each of which defines a specific direction for future work.

The cross-domain evaluation relies on a single independent dataset (DCLDE 2022) of limited duration (~15 minutes), annotated by the first author. While the *N*_*fft*_ ranking is confirmed across 150 models, three experimental arms, and five thresholds, the absolute performance values reported here should not be interpreted as universal benchmarks. Additional crosstest datasets, ideally from different equipment, annotators, geographic regions, and sampling rates — would strengthen the generalizability claim. The framework’s YAML-driven design reduces the overhead of such extensions to the preparation of new annotation files.

The analysis was conducted at a single sampling rate (192 kHz) with a single lightweight CNN architecture. The upsampling-amplification mechanism described in Section 4.2 is sampling-rate dependent: the bin-count per whistle scales linearly with *N*_*fft*_ and inversely with sampling rate, so the optimal *N*_*fft*_ at other sampling rates cannot be assumed to follow a simple scaling rule and should be determined empirically, consistent with the caution in Section 4.4. Similarly, deeper or more expressive architectures may be less sensitive to the spectro-temporal trade-off. As noted in Section 4.4, this means the reported *N*_*fft*_ ranking characterizes the interaction between spectral resolution and this specific architecture, not an architecture-independent property of *N*_*fft*_ itself.

Both directions are directly addressable within the framework by modifying the YAML configuration file, with all downstream stages executing identically. No computational or deployment-cost analysis is presented in this paper; this direction is reserved for future work.

The study is limited to a single species (*T. truncatus*), a single vocalization type (tonal whistles), and a single recording site for training. The mechanism, however, is general: it applies to any scenario in which a frequency-modulated signal occupying a limited number of STFT bins is resized to a fixed CNN input dimension. Whether the specific Pareto point (*N*_*fft*_ = 256) transfers to other tonal species or broader-band vocalizations is an empirical question, but the prediction is clear. The optimal window length should scale with the ratio of vocalization bandwidth to sampling rate.

Despite these constraints, the directional finding is grounded in the physics of the STFT-resize chain: the interaction between spectral bin count, image dimensions, and resize distortion is a geometric property that does not depend on the specific training or test corpus. The consistency of the ranking across 150 models, two overlap conditions, two geometries, and five thresholds provides strong internal replication. Absolute values will change on different data; the conclusion that *N*_*fft*_ has a large, systematic, and direction-consistent effect on cross-domain generalization is expected to hold for tonal cetacean vocalizations at comparable sampling rates.

## 5. Conclusions

This study demonstrates that the FFT window length, a parameter routinely fixed without justification in CNN-based cetacean vocalization detection, has a large and systematic effect on cross-domain generalization, invisible to standard in-domain validation.

1. At 192 kHz, *N*_*fft*_ = 256 is best-performing trade-off observed configuration for binary *T. truncatus* whistle detection: it significantly outperforms longer-window configurations (512, 1024, 2048) in recall and F1, while maintaining a structurally zero false discovery rate across all classification thresholds and experimental conditions. *N*_*fft*_ = 128 achieves comparable recall but fails in specificity, producing false positives in every fold, a failure mode undetectable by in-domain validation.

The *N*_*fft*_ ranking is confirmed across two overlap conditions and five classification thresholds. A supplementary geometry probe (Section 4.4) confirms the ranking further and shows that the magnitude of the geometry effect is itself *N*_*fft*_-dependent, providing direct evidence that the mechanism is the geometric distortion introduced by resizing spectrograms with different native bin structures to a fixed CNN input dimension. The optimal *N*_*fft*_ is not independent of network architecture, however: depth and capacity interact with spectral resolution, and the ranking reported here characterizes one architecture, not a universal rule. Testing *N*_*fft*_ is necessary, but the resulting ranking should be re-verified whenever the architecture changes materially.

Threshold sensitivity analysis serves a dual role in this context: it characterizes operational trade-offs and, more fundamentally, indicates whether a configuration has produced a genuinely separable representation, a diagnostic distinct from, and complementary to, point-estimate accuracy metrics.

Cross-domain evaluation on independently recorded data is necessary to detect the *N*_*fft*_ effect and should be incorporated as standard practice in PAM detector validation. The framework used in this study [18,19] is publicly available and makes such systematic evaluations reproducible with minimal overhead. This study does not resolve the broader multi-parametric space of cetacean CNN detection; it is offered as a template and a prompt for the field to test parameters it currently fixes by convention.

## Acknowledgments

The authors acknowledge the CNR-IRBIM HPC service, with special thanks to Fabrizio Moro and Paolo Scarpini, and the CINECA award under the ISCRA initiative (project id: HP10CL9TFX), for the availability of high-performance computing resources and support.

