## Supplementary material for "Is FFT window length a neutral preprocessing choice in CNN-based dolphin whistle detection? A cross-domain sensitivity analysis" (from another version of this preprint: version 2, 2026-05-15)

#### An Open Reproducible Framework for CNN-Based Cetacean Vocalization Detection in Passive Acoustic Monitoring

**Rocco De Marco**
Institute of Biological Resources and Marine Biotechnology (IRBIM), National Research Council (CNR), Ancona, Italy

---

This document provides the technical depth required to reproduce and extend the experiments, and is self-contained enough to serve as a practitioner reference manual. The complete source code, configuration files, and datasets referenced throughout are available at the DOIs indicated in each section.

---

#### Quick Start — Reproducing the Binary Whistle Detection Experiment

The following ten steps reproduce Experiment A (binary whistle detection, $N_{\text{fft}} = 256$) from a clean system with no prior installation. Estimated time: 15–20 minutes for installation and dataset preparation; training time depends on hardware (GPU recommended).

**Step 1 — Download the toolkit**

Download the versioned source archive from Zenodo:

```
https://doi.org/10.5281/zenodo.19860736
```

**Step 2 — Extract and enter the directory**

```
unzip ai-pam-pipeline-v1.0.zip
cd ai-pam-pipeline
```

**Step 3 — Create a virtual environment**

```
python3 -m venv .venv
source .venv/bin/activate          # Linux / macOS
# .venv\Scripts\activate           # Windows
```

**Step 4 — Install dependencies**

```
pip install -r requirements.txt
```

Core dependencies include TensorFlow, scipy, Pillow, matplotlib, and soundfile. GPU acceleration is available if a CUDA-compatible device is present and the appropriate TensorFlow GPU build is installed (see Section S1).

**Step 5 — Download the Oltremare dataset**

Download the annotated Oltremare dataset from SEANOE:

```
https://doi.org/10.17882/109081
```

**Step 6 — Place audio files**

Copy all `.wav` recording files into the `data/audio/` directory:

```
ai-pam-pipeline/
└── data/
    └── audio/
        ├── 20211120_102109_192.wav
        └── ...
```

**Step 7 — Place label files**

Copy the corresponding Audacity label files (`.txt`) into `data/labels/`. Each label file must share the same stem as its audio file:

```
ai-pam-pipeline/
└── data/
    └── labels/
        ├── 20211120_102109_192.txt
        └── ...
```

**Step 8 — Run preprocessing (Stages 1–4)**

```
python3 ai_pam_preprocessing.py conf/binary_cnn.yaml
```

This command reads the configuration file `conf/binary_cnn.yaml`, processes all paired audio/label files, and exports segmented spectrogram images to `outputs/spectrograms/`. A summary report is printed to the console on completion (see Section S-OUT.1 for a reference output).

**Step 9 — Run training (Stages 5–6)**

```
python3 ai_pam_train.py conf/binary_cnn.yaml -v
```

The `-v` flag enables verbose fold-level progress reporting. Ten-fold cross-validation is executed sequentially; trained models and per-fold metrics are saved to `outputs/models/train/`.

**Step 10 — Inspect results**

Open the training summary report:

```
outputs/report/train/summary_report.md
```

This file contains cross-fold mean ± SD for all metrics, per-fold detail, and a fold-level confusion matrix. Individual `.keras` model files and `_metrics.json` records are available in `outputs/models/` for further analysis or cross-domain evaluation (see Section S8).

---

#### Configuration File Reference — `conf/binary_cnn.yaml`

The entire experiment is defined within a single YAML configuration file, which encompasses all settings for the six pipeline stages. A preview of this configuration is provided below, with further technical details explored in the following sections. This file is the complete reproducibility record for Experiment A at $N_{\mathrm{fft}} = 256$. This YAML files is included in the `conf/` directory of the source archive.

```yaml
# ==============================================================================
# project.yaml — BINARY CNN example
# ==============================================================================
experiment_meta:
  name: "Binary CNN"
  version: "1.0"
  description: "A simple binary CNN example"

# --- Input paths --------------------------------------------------------------
input:
  audio_dir:  "data/audio"       # Directory containing WAV recordings
  label_dir:  "data/labels"      # Directory containing Audacity label files
  audio_glob: "*.wav"
  label_glob: "*.txt"

# --- Output paths -------------------------------------------------------------
report:
  report_dir:     "outputs/report"
  summary_report: false
  full_report:    false
```

```yaml
output:
  spectrogram_dir: "outputs/spectrograms"    # Exported spectrogram images

# --- Pipeline stages ----------------------------------------------------------
pipeline:

  # Stage 1: Audio I/O
  stage_1:
    resample_hz: null       # Keep native sampling rate (192 kHz)
    channel:     null       # Mix to mono
    normalize:   null       # No audio normalization before spectrogram computation

  # Stage 2: Segmentation and windowing
  stage_2:
    tl_step:    0.05        # Timeline discretization step (s)
    as_len:     0.8         # Analysis window length (s)
    as_overlap: 0.5         # Window overlap fraction
    classes:
      C0:                   # Negative class: background noise
        include: ["NOISE"]
        exclude: ["W", "MW"]
        min_duration: 0.15
      C1:                   # Positive class: whistles
        include: ["W", "MW"]
        exclude: ["NOISE", "FB"]
        min_duration: 0.15
    exclude_overlaps:      true    # Apply exclusion zone logic
    center_short_events:   true    # Center events shorter than as_len
    augmentation:
      scroll:
        enabled: false             # Scroll augmentation disabled for Exp. A

  # Stage 3: Spectrogram generation
  stage_3:
    fft_len:      256       # FFT window size (experimental variable in Exp. A)
    fft_overlap:  0.5
    fft_min_freq: 3000      # Frequency crop lower bound (Hz)
    fft_max_freq: 25000     # Frequency crop upper bound (Hz)
    image:
      width_px:  224
      height_px: 224
      dpi:       100
      cmap:      "gray"     # Single-channel grayscale output

  # Stage 4: Image preprocessing
  stage_4:
    image_normalization: true     # Per-image min-max normalization to [0, 1]
    filters:
      mode:     "none"            # No image filters applied in Exp. A
      pipeline: []

  # Stage 5: CNN training
  stage_5:
    model:
      input_shape:  [224, 224, 1]
      architecture: "models/baseline_cnn.py"    # Plug-in model definition
    cross_validation:
      k_folds:               10
      max_samples_per_class: 4700     # Balanced sampling cap
```

```
    training:
      epochs:         100
      batch_size:     32
      learning_rate:  0.0001         # Adam optimizer
      patience:       10             # Early stopping patience (epochs)
      warmup_epochs:  5              # Linear LR warmup
    output:
      model_dir:    "outputs/models"
      dataset_dir: "outputs/spectrograms"    # Must match output.spectrogram_dir

  # Stage 6: Evaluation
  stage_6:
    evaluation:
      threshold: 0.5          # Default sigmoid classification threshold
    crosstest:
      test_dataset_dir:       "crosstest/spectrograms"
      max_samples_per_class: 500
      seed:                   42
      mode:                   "all"    # Per-fold evaluation for Wilcoxon tests
      report_dir:             "crosstest/report"
```

The complete two-level YAML schema — including `master_config.yaml` with all defaults and the full parameter reference table — is described in Section S2.

---

#### S-OUT.1 — Preprocessing Output Overview

The following is the console output produced by `ai_pam_preprocessing.py` on the Oltremare dataset with the `binary_cnn.yaml` configuration ($N_{\mathrm{fft}} = 256$). This output serves as a reference for verifying that a local installation produces consistent results.

```
============================================================
  EXPORT SUMMARY
============================================================
  Mode          : standard
  Dry run       : False
  Output dir    : outputs/spectrograms
  Image mode    : L (cmap='gray')
  Elapsed       : 1068.4s

  Class              Files  Segments   Windows  Exported
  ---------------  -----  ---------  ---------  ---------
  C0                   181       1183       8055       8016
  C1                   204       2590       5681       5665

  Total exported: 13681 spectrogram(s)
```

**Reading the summary table.** Each column has a precise meaning:

- **Files**: number of distinct audio files contributing at least one segment to this class.
- **Segments**: number of annotated events (after `min_duration` filtering) that passed the exclusion zone check and produced at least one valid window.
- **Windows**: total number of sliding-window positions generated from all segments, before boundary clamping.

- **Exported**: windows successfully written to disk after boundary clamping (left-edge ≥ 0, right-edge ≤ file duration). The small difference between Windows and Exported (e.g., 8,055 → 8,016 for C0) reflects windows whose audio span partially exceeds the file boundary and are discarded.

The asymmetry between C0 (8,016) and C1 (5,665) reflects the greater abundance of annotated noise segments relative to whistle segments in the Oltremare dataset. Both classes are subsequently balanced to 4,700 samples per class during training (Section S7.2).

The elapsed time of approximately 18 minutes is representative of an eight-core CPU run on the full 22-hour Oltremare corpus. Processing time scales approximately linearly with total audio duration and is independent of `fft_len` (spectrogram computation is the dominant step).

---

#### S-OUT.2 — Training Output Overview

The following is the condensed log produced by `ai_pam_train.py` for the $N_{\text{fft}} = 256$ binary configuration. The first four lines confirm that the configuration system loaded and validated correctly; the subsequent lines report dataset discovery, class balancing, and per-fold progress.

```
INFO    | lib.ai_pam_config | Loading master config: conf/master_config.yaml
INFO    | lib.ai_pam_config | Loading project config: conf/binary_cnn.yaml
INFO    | lib.ai_pam_config | Validating project config against master...
INFO    | lib.ai_pam_config | Config loaded — experiment: 'Binary CNN'
INFO    | __main__ | Experiment: Binary CNN
INFO    | __main__ | Dataset directory: outputs/spectrograms
INFO    | __main__ |   C0: 8011 images
INFO    | __main__ |   C1: 5664 images
INFO    | __main__ | Classes: ['C0', 'C1'] (binary)
INFO    | __main__ |   C0: 4700 samples selected (from 8011 available)
INFO    | __main__ |   C1: 4700 samples selected (from 5664 available)
INFO    | __main__ | Total dataset: 9400 samples
INFO    | __main__ | Class distribution: {0: 4700, 1: 4700}
INFO    | __main__ | FOLD 01/10
INFO    | __main__ |   Train: 8460, Test: 940
[...]
INFO    | __main__ |   Model saved: outputs/models/fold_01.keras
INFO    | __main__ |   Metrics JSON: outputs/models/fold_01_metrics.json
INFO    | __main__ |   Report MD: report/train/fold_01_report.md
INFO    | __main__ |   Fold 01 done — Accuracy: 0.9915, Macro F1: 0.9915 (1096.6s)
[...]
INFO    | __main__ | FOLD 10/10
INFO    | __main__ |   Train: 8460, Test: 940
[...]
INFO    | __main__ |   Fold 10 done — Accuracy: 0.9777, Macro F1: 0.9777 (1204.6s)

============================================================
  TRAINING COMPLETE — 10 fold(s)
  Accuracy : 0.9844 +/- 0.0041
  Macro F1 : 0.9844 +/- 0.0041
  Models   : outputs/models
  Reports  : report/train
============================================================
```

The dataset discovery block (lines 6–12) shows that the script found 8,011 C0 images and 5,664 C1 images and applied the `max_samples_per_class: 4700` cap to produce a balanced 9,400-sample dataset. This cap is applied before fold partitioning, so each fold receives exactly 8,460 training samples and 940 test samples (a 90/10 split of 9,400).

Each fold produces three output files: a `.keras` model, a `_metrics.json` with the full metric set (accuracy, precision, recall, F1, confusion matrix, per-class metrics), and a `_report.md` summary. The final block reports cross-fold mean ± SD and confirms the output directories. The per-fold elapsed time (roughly 18–20 minutes per fold on CPU) indicates that a full 10-fold run requires approximately three hours without GPU acceleration.

---

This overview has provided a first, end-to-end look at the pipeline from raw audio to cross-domain evaluation results. The three scripts — `ai_pam_preprocessing.py`, `ai_pam_train.py`, and `ai_pam_crosstest.py` — cover the complete experimental workflow, and their console output gives immediate feedback on dataset composition, class balancing, and model performance at each stage. The following sections describe each pipeline stage in full technical detail, including all configuration parameters, implementation notes, and practitioner guidance for adapting the framework to new species, datasets, or experimental designs.

#### S1. Software Installation and Environment

This section covers installation on three target environments: a local workstation (Linux/macOS/Windows).

##### S1.1 System Requirements

| Requirement | Minimum | Recommended |
|---|---|---|
| Python | 3.9 | 3.11 |
| RAM | 8 GB | 16 GB |
| Disk (Oltremare dataset + spectrograms) | 50 GB | 80 GB |
| OS | Linux, macOS, Windows 10+ | Ubuntu 22.04 LTS |
| GPU | — | NVIDIA, CUDA 11.8+ |

GPU acceleration is optional but strongly recommended for Stage 5 training. Preprocessing (Stages 1–4) is CPU-bound and does not benefit from GPU. On a eight-core CPU core, preprocessing the full Oltremare corpus takes approximately 25 minutes; training one 10-fold cross-validation run takes approximately 25–40 minutes for each fold without GPU.

##### S1.2 Installation

**Clone or download the repository.**
The versioned release is available on Zenodo at `https://doi.org/10.5281/zenodo.19860736`. Alternatively, clone the development repository:

```
git clone https://github.com/roccodm/ai-pam-pipeline
cd ai-pam-pipeline
```

**Create and activate a virtual environment.**
A dedicated virtual environment isolates the toolkit dependencies from the system Python installation and is strongly recommended:

```
python3 -m venv .venv
source .venv/bin/activate          # Linux / macOS
# .venv\Scripts\activate           # Windows PowerShell
```

**Install dependencies.**

```
pip install --upgrade pip
pip install -r requirements.txt
```

The `requirements.txt` file is organised into three dependency groups:

| Group | Packages | Required for |
|---|---|---|
| Core | soundfile, scipy, numpy, Pillow, matplotlib, pyyaml | Stages 1–4 |
| Training | tensorflow ≥ 2.12, scikit-learn | Stages 5–6 |
| Dev | pytest, pytest-cov | Test suite |
| Utils | fpdf2 ≥ 2.7 | Spectrogram exporter |

To install all the dependencies:

```
pip install -r requirements.txt
```

**Verify the installation.**

```
python -m pytest tests/ -v
```

All tests should pass. A failed import of `tensorflow` on a preprocessing-only node is expected if the training group was not installed.

**GPU support (optional).**
If a CUDA-compatible GPU is available, install the GPU build of TensorFlow:

```
pip install tensorflow[and-cuda]
```

Verify GPU visibility before running training:

```
python -c "import tensorflow as tf;
print(tf.config.list_physical_devices('GPU'))"
```

An empty list indicates that TensorFlow is running in CPU-only mode.

#### S2. Configuration System

##### S2.1 Two-level YAML Architecture

The *ai-pam-pipeline* framework utilizes a single YAML configuration file designed to provide a comprehensive description of the experimental setup. This file is organized into global configuration blocks—`experiment_meta`, `input`, `output`, and `report`—followed by specific parameter sections dedicated to each of the six pipeline stages, which are detailed in the subsequent sections of this document.

For the studies presented in the main manuscript, the `conf/` directory includes the configuration files for **Experiment A** (`binary_cnn.yaml`) and **Experiment B** (`multiclass_cnn.yaml`). However, the framework is flexible, allowing user-defined configuration files to be stored at any location within the filesystem.

The system relies on a dual-layer configuration logic involving a **Master Configuration** and a **Project Configuration**:

- **Master Configuration (`master-config.yaml`):** Located strictly within the `conf/` directory, this file serves as the global schema. It contains an exhaustive list of all available parameters, each assigned a robust default value. The `master-config.yaml` is extensively commented to serve as a primary reference for the framework's capabilities.
- **User-Defined Configuration:** This file contains experiment-specific overrides. During execution, the framework performs a merge operation between the user-defined file and the master-config. User settings take precedence; default values from the master configuration are only injected when specific keys are absent from the user-defined file.

**Best Practices and Validation:**

For initiating new experiments, the recommended workflow is to clone the `master-config.yaml` into a new project file and modify the parameters according to the experimental requirements.

The framework includes a **validation layer** that monitors the merging process. Any structural inconsistencies—specifically configuration keys present in the user file but missing from the master schema—are detected and flagged as diagnostic errors. Consequently, direct modification of the `master-config.yaml` is strongly discouraged to maintain the integrity of the validation system.

---

##### S2.2 Experiment Reproducibility

To ensure full experimental tracking and reproducibility, the framework automatically exports a `run_config.yaml` file—representing the final result of the merge between the project configuration and the master configuration—into the exported spectrograms folder.

In this way, every generation of spectrograms, whether for training or cross-testing, is clearly linked to its complete preparation settings. This allows for a rigorous verification of parameter consistency by comparing the `run_config.yaml` files from different output directories.

Specifically, to guarantee the integrity of cross-domain evaluations, all spectrogram generation parameters must be identical. By maintaining this "digital twin" of the configuration alongside the data, the *ai-pam-pipeline* effectively prevents training-serving skew and facilitates the validation of results by independent researchers.

##### S2.3 Global parameter blocks

The YAML configuration file features global parameters that, for convenience, are typically placed at the top of the document. These settings govern the general input/output operations, define system paths, and provide bookkeeping metadata to describe the experiment.

##### 1. `experiment_meta`

This block is for human-readable bookkeeping only and is not actively used by any processing stage.

| Key | Type | Description |
|---|---|---|
| `name` | str | Experiment identifier |
| `version` | str | Version tag |
| `description` | str | Free-text description |

These fields can be freely customized by the user. Maintaining accurate and detailed metadata is highly recommended, as it proves particularly advantageous for tracking and cataloging iterative runs with varying parameter combinations.

##### 2. `input`

Locates the raw acoustic data and their corresponding annotations.

| Key | Default | Description |
|---|---|---|
| `audio_dir` | `./data/audio` | Directory containing audio files |
| `label_dir` | `./data/labels` | Directory containing annotation files |
| `audio_glob` | `*.wav` | Glob pattern applied within `audio_dir` |
| `label_glob` | `*.txt` | Glob pattern applied within `label_dir` |

These directories specify the paths to the original acoustic recordings and their corresponding label files. Labels are expected in TSV (Tab-Separated Values) format (`<start_offset>\t<end_offset>\t<label>`), which matches the standard annotation output of PAM analysis tools like Audacity.

Pairing between audio and labels is executed by filename stem: `recording_01.wav` is paired with `recording_01.txt`. Files with no matching counterpart are skipped, and a warning is logged.

**Supported Audio Formats:** The framework supports all audio formats handled by the underlying `soundfile` library. You can verify the available formats in your specific environment using the following Python snippet:

```
>>> import soundfile
>>> soundfile.available_formats()
{'AIFF': 'AIFF (Apple/SGI)', 'AU': 'AU (Sun/NeXT)', 'AVR': 'AVR (Audio Visual
Research)',
 'CAF': 'CAF (Apple Core Audio File)', 'FLAC': 'FLAC (Free Lossless Audio Codec)',
 'HTK': 'HTK (HMM Tool Kit)', 'SVX': 'IFF (Amiga IFF/SVX8/SV16)', 'MAT4': 'MAT4
(GNU Octave 2.0 / Matlab 4.2)',
 'MAT5': 'MAT5 (GNU Octave 2.1 / Matlab 5.0)', 'MPC2K': 'MPC (Akai MPC 2k)',
'MP3': 'MPEG-1/2 Audio',
 'OGG': 'OGG (OGG Container format)', 'PAF': 'PAF (Ensoniq PARIS)', 'PVF': 'PVF
(Portable Voice Format)',
 'RAW': 'RAW (header-less)', 'RF64': 'RF64 (RIFF 64)', 'SD2': 'SD2 (Sound Designer
II)',
```

```
 'SDS': 'SDS (Midi Sample Dump Standard)', 'IRCAM': 'SF (Berkeley/IRCAM/CARL)',
'VOC': 'VOC (Creative Labs)',
 'W64': 'W64 (SoundFoundry WAVE 64)', 'WAV': 'WAV (Microsoft)', 'NIST': 'WAV (NIST
Sphere)',
 'WAVEX': 'WAVEX (Microsoft)', 'WVE': 'WVE (Psion Series 3)', 'XI': 'XI
(FastTracker 2)'}
```

##### 3. `output`

This section defines the target directories for exporting data that will be reused in subsequent pipeline stages or external analyses.

| Key | Default | Description |
|---|---|---|
| `spectrogram_dir` | `./outputs/spectrograms` | Root directory for generated spectrogram images |
| `export_dir` | `null` | Raw audio segment export directory; `null` = disabled |

Specifically, this block allows the user to define paths for the generated 2D spectrograms and/or the isolated raw audio segments. Exporting raw segments is particularly useful for external auditory inspection of the training samples, or for workflows that train 1D-CNN models directly on raw audio waveforms rather than spectrogram images.

Images are written following a strict taxonomic hierarchy: `<spectrogram_dir>/<class_name>/<class>-<label>-<stem>-<offset>.png`. When `export_dir` is set, the corresponding processed audio segment is also written as a WAV file with an identical naming convention.

**Note:** For the training script (`ai_pam_train.py`) to successfully locate the images produced during the preprocessing phase (`ai_pam_preprocessing.py`), the parameter `pipeline.stage_5.output.dataset_dir` must point to the exact same path defined in `output.spectrogram_dir`.

##### 4. `report`

Controls the verbosity and destination of console and file reporting.

| Key | Default | Description |
|---|---|---|
| `report_dir` | `./outputs/reports` | Directory for generated reports |
| `summary_report` | `true` | Write a lightweight per-stage summary |
| `full_report` | `false` | Write an extended report with per-file detail |

---

#### S3. Stage 1 — Audio I/O (`lib/ai_pam_audio.py`)

Stage 1 is responsible for reading audio files, selecting or mixing channels, optionally resampling to a target rate, and applying amplitude normalisation before the signal is passed to the spectrogram generation stage. The three parameters that govern this stage are listed in the table below, followed by a detailed discussion of each.

| Parameter | Default | Description |
|---|---|---|
| `stage_1.resample_hz` | `null` | Target sample rate in Hz; `null` = retain native rate |
| `stage_1.channel` | `null` | Channel selection; `null` = mix to mono; integer = zero-based channel index |
| `stage_1.normalize` | `null` | Amplitude normalisation mode: `null`, `"peak"`, or `"rms"` |

##### S3.1 Memory-efficient seek-based reading

A PAM deployment typically produces continuous recordings of 5 to 60 minutes, often stored as multi-gigabyte WAV or FLAC files. Loading such files entirely into memory for each extraction would be prohibitively expensive and would impose an unnecessary upper bound on the size of the corpus that can be processed on a standard workstation.

`lib/ai_pam_audio.py` avoids this by implementing **seek-based segment extraction**: for each annotated event, the reader opens the file, seeks directly to the frame offset corresponding to the event start time, and reads only the samples spanning `[t_start, t_end]`. The file handle is then closed. Peak RAM consumption at any given moment is therefore proportional to the duration of the longest annotated segment — typically a few seconds — rather than to the total file duration.

##### S3.2 Channel selection

PAM deployments frequently involve multi-channel recordings: stereo hydrophone pairs, towed arrays, or multi-element sensor networks are all common in cetacean monitoring. The `channel` parameter controls how multi-channel files are handled.

| Value | Behaviour |
|---|---|
| `null` | Mix all channels to mono by arithmetic mean (default) |
| `0` | Extract channel 0 (first channel, zero-based index) |
| `1` | Extract channel 1 (second channel) |
| `N` | Extract channel N |

When `null` is set, all channels are averaged sample-by-sample before further processing. This is appropriate when all channels provide independent hydrophone signals of comparable quality and the goal is to maximise SNR through averaging.

When a specific channel index is provided, only that channel is extracted and the others are discarded. This is the correct choice when channels differ in quality, when a specific hydrophone orientation is preferred, or when the file contains channels that are not acoustic recordings (e.g., GPS pulse tracks or trigger signals).

**Example — DCLDE 2022 cross-test dataset.** The DCLDE Oahu recordings used for cross-domain evaluation in the main article are multi-channel towed-array files. The file corresponding to acoustic detection number 46 (15 July 2017, 11:09–11:24 UTC) contains multiple tracks recorded simultaneously by different array elements. Track 3 was selected for annotation and testing based on signal quality, and was extracted by setting `stage_1.channel: 3` in the cross-test preprocessing configuration. Processing only the selected track avoids generating redundant spectrogram datasets from all array elements and keeps the cross-test set directly comparable to the annotated sample set.

##### S3.3 Resampling

| Value | Behaviour |
|---|---|
| `null` | Retain the native sampling rate of the file (recommended) |
| integer Hz | Resample all extracted segments to the specified rate |

Resampling is rarely needed when the training and cross-test datasets share the same hardware and sampling rate, but becomes necessary when integrating recordings from different monitoring systems or when a model trained at one rate must be applied to data collected at another.

Retaining the native rate (`null`) is always preferred when possible, as resampling introduces mild spectral artefacts and increases processing time. If resampling is required, it should be applied identically to all datasets used in the same experimental run and recorded in the `run_config.yaml` alongside the other stage parameters.

##### S3.4 Amplitude normalisation

Amplitude normalisation adjusts the signal level of each extracted segment before spectrogram computation. It is applied per-segment, after any resampling, and operates on the 1D audio waveform.

| Value | Behaviour | Recommended use |
|---|---|---|
| `null` | No normalisation; raw amplitude values passed to STFT | When recordings are already gain-matched or when absolute level information is meaningful |
| `"peak"` | Divide all samples by `max(abs(x))`; output range [−1, 1] | When recording gain varies across sessions or deployments |
| `"rms"` | Scale samples so that `RMS(x) = 1`; preserves waveform shape | When SNR varies systematically across event types or distances |

**Peak normalisation** anchors every segment to the same maximum amplitude regardless of absolute signal level. It is effective when the dynamic range of vocalizations is broad — for example, when the same species is recorded at widely varying distances — and when inter-session gain differences would otherwise produce inconsistent spectrogram brightness. The drawback is that it amplifies quiet segments, including those dominated by noise, to the same peak level as strong vocalization segments, which may reduce the apparent SNR contrast between classes.

**RMS normalisation** scales each segment to unit root-mean-square energy. Because the RMS is an energy measure rather than a peak measure, it is less sensitive to brief transients and more representative of the overall signal power in the window. It is preferred when the goal is to equalize the perceived loudness of events rather than their peak amplitude, and is particularly appropriate for continuous tonal vocalizations such as whistles, where energy is distributed across the duration of the call.

Neither mode alters the waveform shape or the relative frequency content; only the amplitude scale changes. Both are therefore spectrally neutral and do not affect the time–frequency structure of the resulting spectrogram. The choice between the two is dataset-dependent: when in doubt, running the preprocessing with `null` and inspecting the resulting spectrogram brightness distribution across classes is a practical diagnostic step.

**Important:** the normalisation mode must be identical across the training and cross-test preprocessing runs. A mismatch produces a systematic input distribution shift between training and inference, which can cause significant recall degradation without any corresponding change in precision — a failure mode that is difficult to diagnose without examining the `run_config.yaml` files from both runs side by side.

---

#### S4. Stage 2 — Label Parsing, Segmentation, and Windowing (`lib/ai_pam_segment.py`)

Stage 2 is the most structurally complex stage of the pipeline. It reads the Audacity label files produced in Stage 1, maps the annotations onto a discrete timeline, applies exclusion zone logic to remove windows that would contaminate one class with the signal of another, performs binary erosion to trim residual boundary fragments, and finally extracts the set of audio windows that will be rendered as spectrogram images in Stage 3.

The full processing chain for each class is:

```
1. Filter by minimum duration
   |
   └───> 2. Build label matrix
            |
            └───> 3. Apply exclusion zones
                     |
                     └───> 4. Binary erosion
                              |
                              └───> 5. RLE segment extraction
                                       |
                                       └───> 6. Windowing
```

For each processed audio file, the pipeline produces a per-file report and a timeline visualisation showing the annotation state at each step of the chain. An example is shown in Figure S1 below.

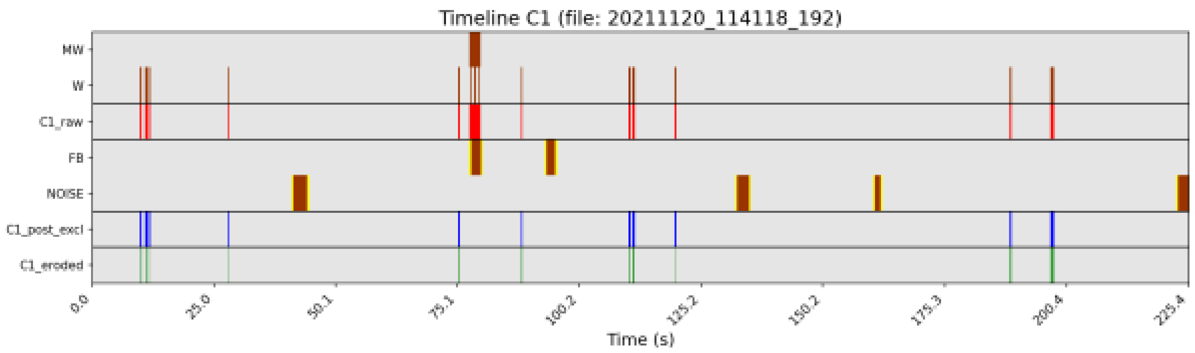

Figure S1: Timeline visualisation for class C1 (whistles) from a 225-second segment of the Oltremare corpus. Each horizontal band shows the annotation state at one step of the Stage 2 processing chain. MW and W rows show the raw Audacity annotations; C1_raw is their union before any filtering; FB and NOISE rows show the exclusion sources; C1_post_excl is the C1 signal after exclusion zone subtraction; C1_eroded is the final sampling domain after binary erosion. Green segments in C1_eroded are the intervals from which spectrogram windows are extracted.

Reading the figure from top to bottom traces the progressive refinement of the usable C1 signal. The MW and W rows show all whistle-type annotations present in the file, some of which co-occur with FB (feeding buzz) or NOISE events. After exclusion zone application (C1_post_excl, blue), the portions of C1 that temporally overlapped with excluded classes are removed, often reducing a long annotation span to a shorter usable fragment or eliminating it entirely. Binary erosion (C1_eroded, green) then contracts each surviving fragment at both ends, removing the thin boundary slices that may still contain mixed-class signal. Only the green segments are eligible for window extraction.

---

##### S4.1 Timeline discretisation

| Parameter | Default | Description |
|---|---|---|
| `stage_2.tl_step` | `0.05` | Frame duration in seconds for the discrete timeline |

All label-file timestamps are converted to integer frame indices by dividing by `tl_step` and rounding. The resulting integer-valued label matrix (one row per annotation class, one column per frame) is the internal representation on which all subsequent operations — exclusion, erosion, and RLE segmentation — are performed. Floating-point accumulation errors in the label file timestamps are absorbed by this discretisation step.

Smaller `tl_step` values produce finer temporal granularity, which improves the precision of exclusion zone boundaries and erosion edges, at the cost of proportionally larger label matrices. Values between 0.02 and 0.10 s are suitable for most PAM applications. For the experiments in the main article, `tl_step: 0.05` s was used throughout, yielding a resolution of one frame per 50 ms.

---

##### S4.2 Class definitions

The `stage_2.classes` key is the only parameter that **must** be defined in the project file; it has no meaningful default in `master_config.yaml`. This design reflects the fact that the class taxonomy is intrinsically experiment-specific: the same audio corpus can be used to train a binary whistle detector, a multiclass vocalization classifier, or a noise characterisation model simply by changing the class definitions in the YAML file, without modifying any code or reprocessing the raw audio.

Each class entry specifies four fields:

| Field | Type | Description |
|---|---|---|
| `include` | list[str] | Audacity label strings treated as positive signal for this class |
| `exclude` | list[str] | Audacity label strings that generate exclusion zones; windows overlapping these labels are discarded |
| `min_duration` | float | Minimum event duration in seconds; shorter events are filtered before discretisation, and the same value is used as the binary erosion kernel radius |
| `augmentation_scroll` | bool\|null | Per-class scroll override: `true` or `false` overrides the global flag; `null` inherits the global setting |

The `include` list defines what counts as a positive event for that class. For C1 (whistles) in the binary experiment, this is `["W", "MW"]`: both single-whistle and multi-whistle annotations contribute positive windows. For C0 (noise), it is `["NOISE"]`.

The `exclude` list defines which co-occurring annotations make a window untrustworthy for that class. For C1, `["NOISE", "FB"]` means that any window whose time span overlaps a NOISE or FB annotation — even partially — is flagged for removal. The symmetric question ("should C0 exclude W and MW?") is also answered in the YAML: the C0 entry lists `["W", "MW", "W?"]` as exclusions, so noise windows that contain even brief whistle activity are discarded from the negative class. This bidirectional exclusion is what gives the segmentation its class-separating power.

---

##### S4.3 Windowing parameters

| Parameter | Default | Description |
|---|---|---|
| `stage_2.as_len` | `0.8` | Analysis window duration in seconds |
| `stage_2.as_overlap` | `0.5` | Fractional overlap between consecutive windows |
| `stage_2.exclude_overlaps` | `true` | Discard any window whose time span overlaps an exclusion zone |
| `stage_2.center_short_events` | `true` | Represent events shorter than `as_len` with a single centred window |

The hop size in seconds is `hop = as_len × (1 − as_overlap)`. For the experimental parameters used in the main article (0.8 s, 50% overlap), the hop is 0.4 s, meaning a new window starts every 400 ms along each eligible segment. The choice of 0.8 s was motivated by the typical duration of *T. truncatus* whistle contours at the Oltremare site; other species or recording scenarios may require different window lengths.

`center_short_events: true` applies when an annotated event is shorter than `as_len`. Rather than generating a window that places the signal at an arbitrary position within the frame — which would depend on the event's absolute timing in the recording — the pipeline generates a single window in which the event is temporally centred. This ensures that all short vocalizations appear at a consistent spatial position within the spectrogram image, which improves training stability and makes the learned features position-invariant with respect to event timing. The default value `true` is recommended for all new experiments.

---

##### S4.4 Exclusion zone logic and binary erosion

The exclusion mechanism operates in two sequential steps, both illustrated in Figure S1.

**Step 1 — Exclusion zone application.** For each class, every label in the `exclude` list is looked up in the label matrix. Its active frames are dilated by a padding margin equal to the hop size (`as_len × (1 − as_overlap)`) applied symmetrically around each annotation boundary. The resulting padded rows are combined into a composite exclusion mask, and any frame in the target class that overlaps this mask is suppressed. The padding margin ensures that windows starting just before or just after an excluded event do not contain residual signal from that event at their edges.

**Step 2 — Binary erosion.** After exclusion, the surviving active spans of the target class may still contain short fragments at their boundaries that are too brief to yield a clean window of the required duration. Binary erosion with a structuring element of length `ceil(min_duration / tl_step)` contracts each contiguous active span at both ends, eliminating these boundary fragments. The operation is implemented via `scipy.ndimage.binary_erosion` on the integer frame representation. Fragments shorter than the erosion kernel are removed entirely; longer spans are trimmed symmetrically.

The two-step design — dilation of excluded regions followed by erosion of the target region — is conservative by construction: it is always preferable to discard a borderline window than to include one that contaminates the class boundary. The per-file timeline visualisation (Figure S1) makes the effect of both steps directly inspectable, allowing the practitioner to verify that the exclusion logic is behaving as expected for a given annotation density.

---

##### S4.5 Per-file reporting

For each audio file processed, Stage 2 writes two output artefacts to the report directory:

- A **Markdown report** (`<stem>_report.md`) summarising the annotation counts, segment counts before and after exclusion and erosion, and the number of windows produced per class.
- A **timeline PNG** (`<stem>_timeline_<class>.png`) showing the label matrix state at each processing step for the target class, in the same format as Figure S1.

These per-file outputs are the primary diagnostic tool for understanding why a particular recording contributed fewer windows than expected. Common causes include a high density of overlapping annotations (producing narrow or zero-length post-exclusion spans), a large proportion of short events filtered by `min_duration`, or file-boundary effects for annotations that begin near the end of a recording.

---

##### S4.6 Scroll augmentation

| Parameter | Default | Description |
|---|---|---|
| `stage_2.augmentation.scroll.enabled` | `false` | Global enable flag |
| `stage_2.augmentation.scroll.scroll_step` | `0.1` | Window advance in seconds between consecutive scroll positions |

When scroll augmentation is enabled, the sliding-window algorithm is applied with step `scroll_step` instead of the standard `as_overlap`-derived hop. For a `scroll_step` of 0.1 s, a 2-second annotated event yields approximately 12–15 windows instead of the 4–5 that standard windowing would produce, each capturing the vocalization at a slightly different temporal position within the frame. This diversity of spatial positions acts as an implicit positional augmentation, making the trained model less sensitive to the precise temporal alignment of a vocalization within the analysis window.

Scroll augmentation is activated globally via `augmentation.scroll.enabled: true` and can be overridden on a per-class basis via `classes.<name>.augmentation_scroll`. This per-class override is useful when augmentation is needed only for a minority class with few annotated events — for example, in Experiment B of the main article, scroll augmentation was enabled selectively for C4 (feeding buzzes), which had fewer than 600 original annotated segments, while remaining disabled for the other classes. When `center_short_events: true` and scroll is simultaneously enabled, scroll takes precedence for events shorter than `as_len`.

---

#### S5. Stage 3 — Spectrogram Generation (`lib/ai_pam_spectro.py`)

Stage 3 converts each audio window produced by Stage 2 into a spectrogram image ready for CNN input or visual inspection. The conversion follows three steps: computation of the Short-Time Fourier Transform (STFT) using a Hann window, frequency-domain cropping to the bandwidth of interest, and rendering to a fixed-size image via a shared `matplotlib` Figure exported through PIL. The mathematical formulation and the physical interpretation of the time–frequency trade-off are discussed in Sections 2.4 and 4.1 of the main article respectively; this section documents the YAML parameters and implementation details relevant for reproducing or extending the experiments.

The effect of `fft_len` on the rendered spectrogram is directly visible in Figure 5 of the main article, which shows the same three whistle events rendered at $N_{\mathrm{fft}} \in \{256,512,1024\}$: coarser spectral bins produce wider, higher-contrast FM traces after bilinear upsampling to the fixed 224×224 pixel grid, while finer bins yield thinner, more acoustically detailed contours at reduced local contrast.

---

##### S5.1 FFT parameters

| Parameter | Default | Description |
| --- | --- | --- |
| `stage_3.fft_len` | `512` | FFT window length in samples; the experimental variable in Experiment A |
| `stage_3.fft_overlap` | `0.5` | Fractional overlap between consecutive FFT frames (range 0–1) |
| `stage_3.fft_min_freq` | `3000` | Lower frequency bound for spectral cropping (Hz) |
| `stage_3.fft_max_freq` | `25000` | Upper frequency bound for spectral cropping (Hz) |

The hop size in samples is `fft_len × (1 − fft_overlap)`. The magnitude spectrogram is converted to a dB scale via $20 \log_{10}(|X| + \varepsilon)$ with $\varepsilon = 1\times10^{-10}$ to prevent logarithm of zero. Frequency cropping is performed by discarding STFT magnitude bins outside `[fft_min_freq, fft_max_freq]` before rendering, which both reduces input dimensionality and focuses the CNN on the biologically relevant bandwidth. For *T. truncatus* whistle detection, 3–25 kHz encompasses the typical FM contour range while excluding the dominant low-frequency environmental noise floor.

---

##### S5.2 Image output parameters

| Parameter | Default | Description |
| --- | --- | --- |
| `stage_3.image.width_px` | `224` | Output image width in pixels |
| `stage_3.image.height_px` | `224` | Output image height in pixels |
| `stage_3.image.dpi` | `100` | Dots per inch for the rendered figure |
| `stage_3.image.cmap` | `"gray"` | Matplotlib colormap name |
| `stage_3.image.show_axis` | `false` | Include frequency and time axis labels |

All spectrogram images are rendered using a singleton `matplotlib` Figure/Axes instance initialised once at startup, avoiding repeated figure allocation overhead during batch export. The rendering pipeline is: STFT computation → frequency slicing → `imshow` on the shared Axes → `fig.canvas.draw()` → RGBA buffer → PIL Image conversion.

**Colormap and channel mode.** When `cmap` is set to `"gray"` or any of its variants (`"Greys"`, `"gray_r"`, `"Greys_r"`), the PIL output mode is `"L"` (8-bit single-channel grayscale), which is the correct format for a `[H, W, 1]` CNN input. Any other colormap — `"plasma"`, `"viridis"`, `"jet"`, etc. — produces an `"RGB"` three-channel output. The output mode is determined once at initialisation and propagates consistently through Stage 4 filters and Stage 5 training, preventing silent channel-count mismatches. For the experiments in the main article, `cmap: "gray"` was used throughout.

**Axis labels.** When `show_axis: true`, the image includes frequency (kHz) and time (s) tick marks and axis labels with automatic scaling derived from the STFT parameters. This mode is intended for visual inspection and reporting, not for CNN training, as the axis border area reduces the usable spectrogram region within the fixed pixel budget.

**Large-format images for reporting.** The `width_px`, `height_px`, and `dpi` parameters can be set to any value independently of the CNN input size. A typical high-resolution report image uses `width_px: 1200`, `height_px: 1200`, `dpi: 150`, `show_axis: true`, and a colour colormap, producing a publication-quality spectrogram with labelled axes that is visually distinct from the compact training images. This configuration is useful for generating figures for papers or monitoring dashboards without modifying the training pipeline configuration.

**Important:** the `stage_3` parameters — particularly `fft_len`, `fft_min_freq`, `fft_max_freq`, `width_px`, `height_px`, and `image_normalization` — must be identical across the training and cross-test preprocessing runs. Any mismatch produces a systematic input distribution shift that degrades cross-domain performance, often manifesting as a recall collapse that is not accompanied by a change in precision and is therefore difficult to diagnose without comparing the `run_config.yaml` files from both directories side by side.

---

#### S6. Stage 4 — Image Preprocessing (`lib/ai_pam_filters.py`)

Stage 4 applies two independent operations to each rendered spectrogram image before it is written to disk: per-image normalisation and an optional OpenCV filter pipeline. Both are controlled entirely via YAML and require no code changes to configure. The stage is optional in the sense that both operations can be disabled (normalization set to `false`, mode set to `none`), but normalisation is enabled by default and its use is recommended for all new experiments.

A complete reference for all supported filters, including YAML schema, parameter descriptions, pipeline modes, compound recipes, and error diagnostics, is provided in `FILTER_REFERENCE.md` at the root of the repository. This section summarises the key concepts and provides two representative examples.

---

##### S6.1 Normalisation

| Parameter | Default | Description |
|---|---|---|
| `stage_4.image_normalization` | `true` | Apply min–max normalisation to pixel values |

When enabled, pixel intensities are linearly rescaled to [0, 255] independently for each image, using the per-image minimum and maximum as the rescaling bounds. This compensates for amplitude variability across recording sessions, hydrophone sensitivities, and distance-to-source variation, ensuring that the CNN receives inputs with a consistent dynamic range regardless of the absolute signal level in the original recording.

For uniform images (min == max, which can occur for near-silent segments), the image is returned unchanged to avoid division by zero.

**This setting must be identical in the training and cross-test preprocessing runs.** A mismatch introduces a systematic input distribution shift that typically manifests as a recall collapse while precision remains unaffected, making the cause difficult to diagnose without comparing the `run_config.yaml` files from both directories.

---

##### S6.2 Filter pipeline

| Parameter | Default | Description |
|---|---|---|
| `stage_4.filters.mode` | `"none"` | Filter application mode |
| `stage_4.filters.pipeline` | `[]` | Ordered list of filter specifications |

**Filter modes.**

| Value | Behaviour |
|---|---|
| `"none"` | No filtering; images passed through normalisation only |
| `"sequential"` | Filters applied in order; output of each feeds the next |
| `"mix"` | Each filter applied independently to the original; results blended as a weighted sum clipped to [0, 255] |

The filter module is built around a generic dispatch mechanism: any `cv2` function is accessible by name without a hardcoded filter list. Two calling conventions are supported — direct filters (the image is passed as the first positional argument) and factory filters (a constructor creates an object, then a method on that object processes the image). The distinction is controlled by the `factory` flag in each pipeline entry.

Each entry in `pipeline` accepts the following fields:

| Field | Type | Default | Description |
|---|---|---|---|
| `name` | str | required | `cv2` function or constructor name |
| `args` | list | `[]` | Positional arguments (after the image for direct filters) |
| `kwargs` | dict | `{}` | Keyword arguments |
| `factory` | bool | `false` | `true` for constructor-based filters (e.g. `createCLAHE`) |
| `apply_method` | str | `"apply"` | Method name on factory |

| Field | Type | Default | Description |
|---|---|---|---|
| | | | output |
| `input_convert` | str\|null | `null` | Force PIL mode conversion before this filter (e.g. `"L"`) |
| `weight` | float | `1.0` | Blending weight for `mix` mode |

All errors produce a diagnostic block containing the filter name, args, kwargs, and input image shape before raising, allowing diagnosis without a debugger. Common error causes and their fixes are documented in `FILTER_REFERENCE.md §7`.

---

##### S6.3 Representative examples

The following two examples illustrate the most common use cases for PAM spectrogram preprocessing. Both are drawn from the recipes in `FILTER_REFERENCE.md`; additional examples covering smoothing, morphological operators, and polarity inversion are provided there.

**Example 1 — CLAHE (adaptive contrast enhancement).**
CLAHE (Contrast Limited Adaptive Histogram Equalization) improves local contrast by equalising the pixel intensity histogram within small image tiles, with a configurable clip limit that prevents excessive noise amplification. It is effective on spectrograms with non-uniform background noise, where it makes faint whistle contours more visually prominent without saturating regions of high signal intensity. CLAHE requires single-channel grayscale input; the `input_convert: "L"` field ensures this regardless of the global colormap setting.

```
stage_4:
  image_normalization: true
  filters:
    mode: sequential
    pipeline:
      - name: createCLAHE
        factory: true
        input_convert: "L"
        kwargs:
          clipLimit: 2.0
          tileGridSize: [8, 8]
        apply_method: apply
```

**Example 2 — CLAHE + Sobel sequential chain (whistle edge enhancement).**
This two-step chain first enhances local contrast with CLAHE, then applies a horizontal Sobel operator (first-order derivative in the frequency axis direction) to accentuate the boundary of FM contours. The sequential mode means the Sobel filter receives the CLAHE-processed image, not the original. This combination is the recipe shown in Figure 3 of the main article (panels b and c). The `ddepth` value `-1` preserves the input bit depth; using `cv2.CV_64F` (value `6`) instead allows negative gradients, which are then normalised to uint8 by the module automatically.

```yaml
stage_4:
  image_normalization: true
  filters:
    mode: sequential
    pipeline:
      - name: createCLAHE
        factory: true
        input_convert: "L"
        kwargs:
          clipLimit: 2.0
          tileGridSize: [8, 8]
        apply_method: apply
      - name: Sobel
        args: [-1, 1, 0]       # ddepth=-1, dx=1 (frequency axis), dy=0
        kwargs:
          ksize: 3
```

For a full filter reference including mix-mode recipes, the `bilateralFilter`, `medianBlur`, `Laplacian`, `Scharr`, `equalizeHist`, and `GaussianBlur` filters, and guidance on calling filters directly from Python without YAML, see `FILTER_REFERENCE.md` in the repository root.

---

#### S7. Stage 5 — CNN Training (`ai_pam_train.py`)

Stage 5 assembles the spectrogram dataset from the images produced by Stage 4, runs stratified k-fold cross-validation, and writes one trained model per fold together with its performance metrics and diagnostic outputs. The training loop is fully decoupled from the model architecture: the CNN is loaded dynamically from the Python file specified in the YAML, allowing any Keras-compatible architecture to be substituted without modifying the training code.

---

##### S7.1 Global training seed

| Parameter | Default | Description |
|---|---|---|
| `stage_5.seed` | `42` | Controls fold partitioning, weight initialisation, and sample shuffling |

The seed is re-applied at the start of each fold to ensure that weight initialisation is independently reproducible across folds. Combined with the `run_config.yaml` record, this guarantees that any reported result can be exactly regenerated from the original spectrogram dataset.

---

##### S7.2 Model architecture

| Parameter | Default | Description |
|---|---|---|
| `stage_5.model.input_shape` | `[224, 224, 1]` | CNN input dimensions `[height, width, channels]`; must match Stage 3 image settings |

| Parameter | Default | Description |
|---|---|---|
| `stage_5.model.architecture` | `models/baseline_cnn.py` | Path to a Python file defining `build_model(input_shape, n_classes, kernel_size)` |
| `stage_5.model.kernel_size` | `[3, 3]` | Convolutional kernel size applied to all blocks |

The architecture plug-in convention requires only that the target Python file expose a `build_model(input_shape, n_classes, kernel_size)` function returning an uncompiled `tf.keras.Model`. The training loop calls this function, compiles the model, and manages the rest. Binary models use a sigmoid output unit with binary cross-entropy loss; multiclass models use a softmax output layer with categorical cross-entropy, selected automatically based on the number of classes defined in Stage 2.

**Baseline architecture (`models/baseline_cnn.py`).** The default model used in both experiments of the main article is a lightweight Sequential CNN designed for deployment on resource-constrained hardware. It consists of three convolutional blocks with an increasing number of filters (32 → 64 → 128), each followed by ReLU activation and 2×2 max-pooling, then a fully connected head with 128 units and the appropriate output activation. The kernel size is configurable via YAML and is applied uniformly to all convolutional layers.

```
Input  [224 × 224 × 1]
  Conv2D(32,  3×3, ReLU) → MaxPool(2×2)
  Conv2D(64,  3×3, ReLU) → MaxPool(2×2)
  Conv2D(128, 3×3, ReLU) → MaxPool(2×2)
  Flatten
  Dense(128, ReLU)
  Dense(1, sigmoid)       ← binary
  Dense(N, softmax)       ← multiclass (N classes)
```

To use a custom architecture, copy `baseline_cnn.py`, edit `build_model`, and point the YAML to the new file:

```yaml
stage_5:
  model:
    architecture: models/my_architecture.py
```

No other configuration change is required.

---

##### S7.3 Cross-validation

| Parameter | Default | Description |
|---|---|---|
| `stage_5.cross_validation.k_folds` | `10` | Number of folds |
| `stage_5.cross_validation.stratified` | `true` | Preserve class distribution in each fold |
| `stage_5.cross_validation.max_samples` | `null` | Cap per class before folding; `null` |

| Parameter | Default | Description |
|---|---|---|
| `_per_class` | | = use all available samples |

Stratified partitioning is recommended whenever class frequencies differ. For the balanced binary design of Experiment A — where C0 and C1 are capped to the same number of samples before folding — stratified and standard k-fold are equivalent. For Experiment B (five-class, unequal support), stratified partitioning ensures that each fold contains a representative proportion of all classes, including the minority class C2.

---

##### S7.4 Training hyperparameters

| Parameter | Default | Description |
|---|---|---|
| `stage_5.training.epochs` | `100` | Maximum training epochs per fold |
| `stage_5.training.batch_size` | `32` | Mini-batch size |
| `stage_5.training.learning_rate` | `0.0001` | Initial Adam optimiser learning rate |
| `stage_5.training.patience` | `10` | Early stopping patience in epochs |
| `stage_5.training.warmup_epochs` | `5` | Epochs during which early stopping is suppressed |
| `stage_5.training.restore_best_weights` | `true` | Restore weights from the lowest-validation-loss epoch on stopping |

Training uses the Adam optimiser with fixed learning rate. Early stopping monitors validation loss with a minimum improvement threshold of $1\times10^{-4}$; if no improvement is observed for `patience` consecutive epochs, training halts and the best weights are restored. The `warmup_epochs` parameter suppresses early stopping during the first epochs of training, preventing premature termination during the initial loss plateau that is common in multiclass softmax configurations.

---

##### S7.5 Training outputs

| Parameter | Default | Description |
|---|---|---|
| `stage_5.output.model_dir` | `./outputs/models` | Directory for model files and metric records |
| `stage_5.output.dataset_dir` | `./outputs/spectrograms` | Spectrogram directory to load; must match `output.spectrogram_dir` |

For each fold, the following files are written to `model_dir`:

- **`fold_NN.keras`** — the trained model at the best-validation-loss epoch, in Keras native format. Used directly by `ai_pam_crosstest.py` for cross-domain evaluation.

- `fold_NN_metrics.json` — the complete metric set for the held-out fold: accuracy, precision, recall, F1, macro F1, weighted F1, confusion matrix, and per-class precision/recall/F1.
- `fold_NN_curves.png` — training and validation loss and accuracy curves across epochs, with a vertical marker at the best epoch selected by early stopping (Figure S2 below).
- `fold_NN_report.md` — a human-readable summary combining the metrics and a text rendering of the confusion matrix.

After all folds complete, a `summary_report.md` is written with cross-fold mean ± SD for all aggregate metrics.

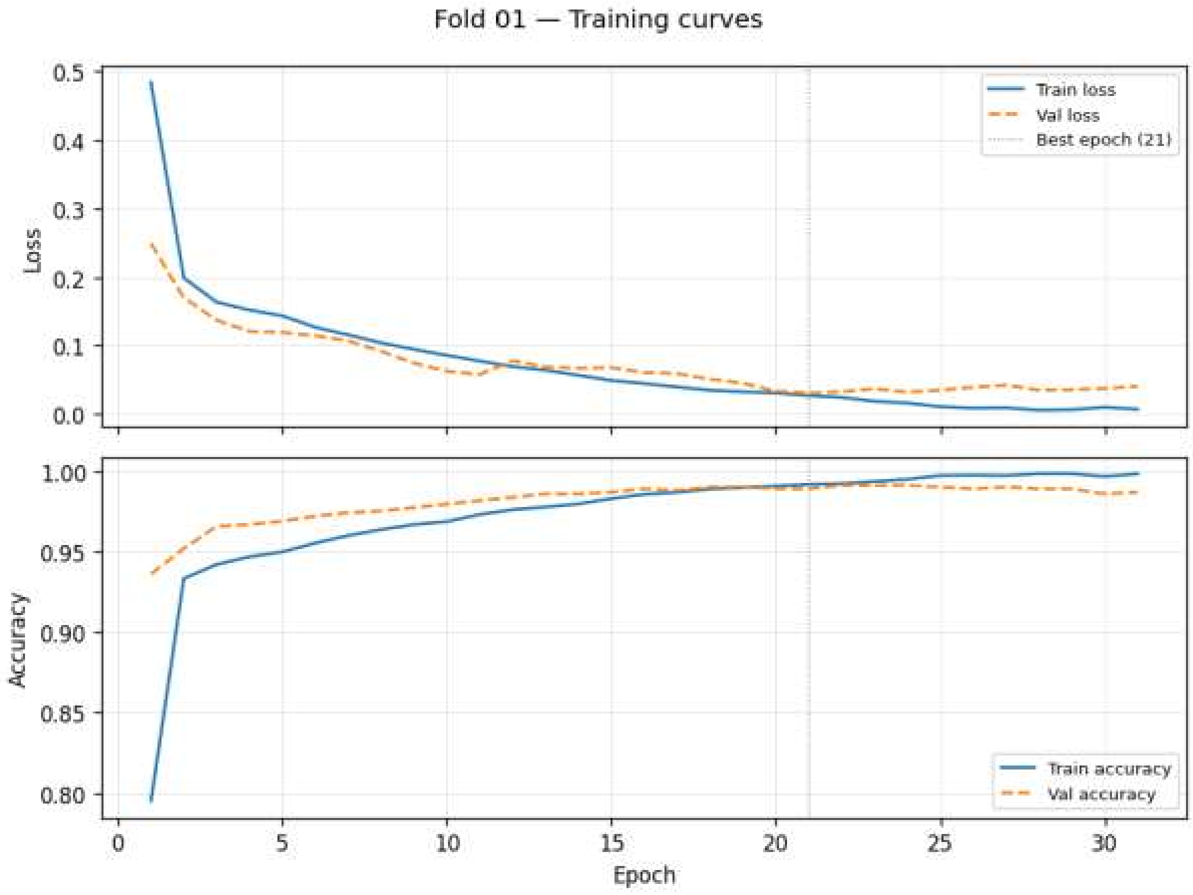

Figure S2: Training curves for fold 01 of the N_fft=256 binary configuration (Experiment A). Upper panel: training loss (blue solid) and validation loss (orange dashed). Lower panel: training accuracy and validation accuracy. The vertical dotted line marks the best epoch (epoch 21) at which weights were restored by early stopping. The gap between training and validation loss after epoch 21 reflects continued training beyond the optimal point; the restored weights correspond to the minimum of the validation loss curve.

The training curve is the primary diagnostic for assessing whether a model has converged cleanly. A well-behaved run shows validation loss decreasing monotonically before the best epoch with no divergence between training and validation accuracy, as in Figure S2. Signs of overfitting — a validation loss that stops decreasing or begins to rise while training loss continues to fall — are addressed by reducing `max_samples_per_class`, increasing `patience`, or applying stronger regularisation in the model architecture. Underfitting (both curves plateau at a high loss value) suggests an insufficient learning rate, too few epochs, or a mismatch between `input_shape` and the actual image dimensions.

#### S8. Stage 6 — Cross-Domain Evaluation (`ai_pam_crosstest.py`)

Stage 6 evaluates the fold models trained in Stage 5 against an independent spectrogram dataset that was not used at any point during training. Cross-domain evaluation is the most demanding test of a PAM model because it exposes the gap between in-domain generalisation — measured by cross-validation on the training corpus — and deployment performance on unseen equipment, sites, and acoustic conditions.

The workflow for Stage 6 consists of three sequential steps: annotating the external audio, generating its spectrogram dataset via a dedicated YAML configuration, and running `ai_pam_crosstest.py` against the trained models.

---

##### S8.1 Step 1 — Annotating the external dataset

The external audio must be annotated following the same labelling protocol used for the training corpus (Section 2.2 of the main article and Section S4.2 of this document). Labels are created in Audacity and exported as TSV files using the same stem-matching convention: `recording.flac` → `recording.txt`.

For the DCLDE 2022 evaluation reported in the main article, track 3 of acoustic detection number 46 was annotated independently by the author, yielding 458 whistle segments (label W) and the corresponding non-whistle spans (label NOISE). The annotation was performed without access to the training labels, ensuring a blind cross-domain test. The resulting label file follows the same format as the Oltremare labels and is processed identically by the pipeline.

The key requirement at this step is **label vocabulary consistency**: the `include` and `exclude` lists in the cross-test YAML must use the same label strings that appear in the external annotation file. If the external annotator uses different abbreviations (e.g. `W1` instead of `W`), the YAML `include` list must be updated accordingly.

---

##### S8.2 Step 2 — Generating the cross-test spectrogram dataset

The cross-test spectrogram dataset is produced by running `ai_pam_preprocessing.py` with a **dedicated configuration file** that differs from the training YAML only in the `input`, `output`, and `report` paths. All Stage 3 and Stage 4 parameters — `fft_len`, `fft_overlap`, `fft_min_freq`, `fft_max_freq`, `image.width_px`, `image.height_px`, `image_normalization` — must be identical to the training configuration. Any mismatch produces a systematic input distribution shift that degrades cross-domain performance in ways that are difficult to diagnose.

The configuration used for the DCLDE 2022 cross-test in the main article is reproduced below:

```yaml
experiment_meta:
  name: Binary CNN DCLDE Crosstest
  version: '1.0'
  description: DCLDE Crosstest

input:
  audio_dir:  data/DCLDE/audio    # directory containing the external WAV/FLAC
files
```

```yaml
  label_dir:  data/DCLDE/labels   # directory containing the external label files
  audio_glob: '*.flac'
  label_glob: '*.txt'

output:
  spectrogram_dir: outputs/crosstest/spectrograms

report:
  report_dir: outputs/crosstest/report

pipeline:
  stage_1:
    resample_hz: null
    channel:     null
    normalize:   null             # must match the training configuration

  stage_2:
    tl_step:    0.05
    classes:
      C0:
        include: ["NOISE"]
        exclude: ["W", "MW", "W?"]
        min_duration: 0.15
      C1:
        include: ["W", "MW"]
        exclude: ["NOISE", "FB"]
        min_duration: 0.15
    as_len:     0.8
    as_overlap: 0.5
    exclude_overlaps:    true
    center_short_events: true
    augmentation:
      scroll:
        enabled:     false
        scroll_step: 0.1

  stage_3:
    fft_len:      256             # must match the training configuration
    fft_overlap:  0.5
    fft_min_freq: 3000
    fft_max_freq: 25000
    image:
      width_px:  224
      height_px: 224
      dpi:       100
      cmap:      gray
      show_axis: false

  stage_4:
    image_normalization: true     # must match the training configuration
    filters:
      mode:     none
      pipeline: []
```

Run preprocessing on the external dataset:

```
python ai_pam_preprocessing.py conf/binary_cnn_dclde_crosstest.yaml
```

After completion, verify parameter consistency by comparing the two `run_config.yaml` files:

```
diff outputs/spectrograms/run_config.yaml \
     outputs/crosstest/spectrograms/run_config.yaml
```

The only expected differences are the `input` paths, `output` paths, and `_run_info.timestamp_utc`. Any difference in Stage 3 or Stage 4 parameters indicates a misconfiguration that must be corrected before proceeding.

---

##### S8.3 Step 3 — Running the cross-test

With the cross-test spectrograms in place, cross-domain evaluation is launched with:

```
python ai_pam_crosstest.py conf/binary_cnn_256.yaml
```

The script reads the `stage_6.crosstest` block from the training configuration to locate the model directory and test dataset directory, loads all fold models, and evaluates them on the test set according to the specified mode.

**Reference output for the DCLDE 2022 evaluation, $N_{\text{fft}} = 256$, `mode: all`:**

```
INFO    | lib.ai_pam_config | Config loaded — experiment: 'Binary CNN'
INFO    | __main__ | Cross-test mode    : all
INFO    | __main__ | Classes            : ['C0', 'C1']
INFO    | __main__ | Test dataset dir   : crosstest/spectrograms
INFO    | __main__ | Model dir          : outputs/models
INFO    | __main__ | Threshold          : 0.5
INFO    | __main__ | Discovering test dataset ...
INFO    | __main__ |   C0: 500 image(s)
INFO    | __main__ |   C1: 500 image(s)
INFO    | __main__ | Total test images  : 1000
INFO    | __main__ | Found 10 model file(s).
INFO    | __main__ | Loading test images ...
=======================================================
INFO    | __main__ | Fold 01/10  — fold_01.keras
INFO    | __main__ |   Accuracy: 0.8910  Macro F1: 0.8897  (2.3s)
[...]
INFO    | __main__ | Fold 10/10  — fold_10.keras
INFO    | __main__ |   Accuracy: 0.8840  Macro F1: 0.8824  (1.9s)
=======================================================
  CROSS-TEST COMPLETE — 10 fold model(s)
  Accuracy : 0.8760 +/- 0.0092
  Macro F1 : 0.8740 +/- 0.0097
  Report   : crosstest/report/crosstest
=======================================================
```

The header block confirms the evaluation configuration before loading any model. The per-fold lines (one per fold) report accuracy and macro F1 on the 1,000-image balanced test set; their elapsed times (approximately 2 seconds per fold) confirm that Stage 6 is inference-only and does not involve retraining. The summary block reports the cross-fold mean ± SD, which correspond directly to the values in Table 5 of the main article. The 10 per-fold F1 values visible in the full log (not truncated above) are the paired observations used in the Wilcoxon signed-rank tests of Table 6.

---

##### S8.4 Classification threshold

| Parameter | Default | Description |
|---|---|---|
| `stage_6.evaluation.threshold` | `0.5` | Sigmoid decision threshold for binary classification |

For binary models, a sample is assigned to the positive class (C1, whistle present) if the sigmoid output exceeds `threshold`, and to the negative class otherwise. Adjusting this value shifts the precision–recall trade-off: lower values increase recall at the cost of more false positives; higher values increase precision at the cost of missed detections. The threshold sensitivity analysis reported in Table 7 of the main article was produced by running `ai_pam_crosstest.py` five times with `threshold` set to {0.1, 0.3, 0.5, 0.7, 0.9} in turn, without retraining.

For multiclass models, the predicted class is the argmax of the softmax output vector and the `threshold` parameter has no effect.

---

##### S8.5 Evaluation modes and additional parameters

| Parameter | Default | Description |
|---|---|---|
| `stage_6.crosstest.test_dataset_dir` | `null` | Directory containing cross-test spectrogram images |
| `stage_6.crosstest.max_samples_per_class` | `null` | Cap per class; `null` = use all available images |
| `stage_6.crosstest.seed` | `42` | Random seed for balanced sampling |
| `stage_6.crosstest.mode` | `"all"` | Evaluation mode |
| `stage_6.crosstest.report_dir` | `null` | Output directory; `null` = `<report.report_dir>/crosstest/` |
| `stage_6.crosstest.save_errors` | `false` | Write a TSV of misclassified filenames |

**Evaluation modes.**

| Value | Behaviour | Use case |
|---|---|---|
| `"all"` | Each fold model evaluated independently; 10 metric sets produced | Required input for paired Wilcoxon tests |
| `"ensemble"` | Sigmoid outputs averaged across all fold models before thresholding | Single stable summary metric for reporting |
| `"best"` | Single fold model with highest macro-F1 on the test set | Optimistic upper bound; useful for debugging |

`"all"` and `"ensemble"` produce complementary views of the same model set and can be run sequentially without retraining. In the main article, `"all"` was used to generate the per-fold F1 values for the Wilcoxon tests (Table 6), while `"ensemble"` was used for the cross-domain performance summary (Tables 5 and 7).

When `save_errors: true`, an additional TSV file is written listing each misclassified image with its filename, true class, and predicted class. This file is useful for identifying systematic error patterns — for example, whether false negatives are concentrated in short whistles, specific frequency ranges, or particular recording files.

---

#### S9. Statistical Analysis Details

Once both the training phase (which includes in-domain testing for each fold) and the cross-domain evaluation on an external dataset (such as the DCLDE dataset used in this study) are completed, a `.csv` file can be prepared as follows. This file specifies the dataset name (Oltremare or DCLDE), the $N_{\mathrm{fft}}$ value used in the experiment, and the fold-specific performance metrics (Accuracy, Precision, Recall, F1 Score, and FDR). A truncated version of the CSV containing the results from the experiments conducted is provided below.

| Dataset | NFFT | Fold | Accuracy | Precision | Recall | F1 Score | FDR |
|---|---|---|---|---|---|---|---|
| **Oltremare** | 256 | 1 | 0.9915 | 0.9936 | 0.9894 | 0.9915 | 0.0064 |
| … | | | | | | | |
| Oltremare | 256 | 10 | 0.9777 | 0.9828 | 0.9723 | 0.9777 | 0.0172 |
| **Oltremare** | 512 | 1 | 0.9862 | 0.9893 | 0.9830 | 0.9862 | 0.0107 |
| … | | | | | | | |
| Oltremare | 512 | 10 | 0.9819 | 0.9829 | 0.9809 | 0.9819 | 0.0171 |
| **Oltremare** | 1024 | 1 | 0.9883 | 0.9852 | 0.9915 | 0.9883 | 0.0148 |
| … | | | | | | | |
| Oltremare | 1024 | 10 | 0.9862 | 0.9831 | 0.9894 | 0.9862 | 0.0169 |
| **DCLDE** | 256 | 1 | 0.8910 | 1.0000 | 0.7820 | 0.8897 | 0.0000 |
| … | | | | | | | |
| DCLDE | 256 | 10 | 0.8840 | 1.0000 | 0.7680 | 0.8824 | 0.0000 |
| **DCLDE** | 512 | 1 | 0.8080 | 1.0000 | 0.6160 | 0.8007 | 0.0000 |

| Dataset | NFFT | Fold | Accuracy | Precision | Recall | F1 Score | FDR |
|---|---|---|---|---|---|---|---|
| … | | | | | | | |
| DCLDE | 512 | 10 | 0.7690 | 1.0000 | 0.5380 | 0.7560 | 0.0000 |
| **DCLDE** | 1024 | 1 | 0.8180 | 1.0000 | 0.6360 | 0.8118 | 0.0000 |
| … | | | | | | | |
| DCLDE | 1024 | 10 | 0.7540 | 1.0000 | 0.5080 | 0.7382 | 0.0000 |

The following simplified R program allows for the construction of pairwise tests within the same dataset, comparing $N_{\text{fft}}$ 256 against 512 and 1024, and finally 512 against 1024. To use the script, simply ensure it loads the correct CSV file, adjusting the `sep` (separator) and `dec` (decimal) parameters if necessary to match your local file format.

```
df <- read.csv("results_nfft.csv")

for (ds in c("Oltremare", "DCLDE")) {
  for (metric in c("f1", "recall")) {
    cat("\n=== DATASET:", ds, "| METRIC:", toupper(metric), "===\n")
    pairs <- list(c(256, 512), c(512, 1024), c(256, 1024))
    for (p in pairs) {
      x <- df[df$dataset == ds & df$nfft == p[1], metric]
      y <- df[df$dataset == ds & df$nfft == p[2], metric]
      wt <- wilcox.test(x, y, paired = TRUE, exact = FALSE)
      n <- sum(x != y)
      mu_w  <- n * (n + 1) / 4
      sig_w <- sqrt(n * (n + 1) * (2 * n + 1) / 24)
      z_val <- (wt$statistic - mu_w) / sig_w
      r_eff <- abs(z_val) / sqrt(n)
      cat(sprintf("%4s vs %4s | W=%2.0f | p-value=%.5f | r=%.3f\n",
                  p[1], p[2], wt$statistic, wt$p.value, r_eff))
    }
  }
}
```

The execution results are presented below:

```
=== DATASET: Oltremare | METRIC: F1 ===
 256 vs  512 | W=44 | p-value=0.10247 | r=0.532
 512 vs 1024 | W=32 | p-value=0.68250 | r=0.145
 256 vs 1024 | W=39 | p-value=0.05780 | r=0.652

=== DATASET: Oltremare | METRIC: RECALL ===
 256 vs  512 | W=20 | p-value=0.83363 | r=0.099
 512 vs 1024 | W=32 | p-value=0.64625 | r=0.161
 256 vs 1024 | W=33 | p-value=0.23573 | r=0.415

=== DATASET: DCLDE | METRIC: F1 ===
 256 vs  512 | W=55 | p-value=0.00592 | r=0.886
 512 vs 1024 | W=42 | p-value=0.15358 | r=0.467
 256 vs 1024 | W=55 | p-value=0.00592 | r=0.886
```

```
=== DATASET: DCLDE | METRIC: RECALL ===
 256 vs  512 | W=55 | p-value=0.00589 | r=0.886
 512 vs 1024 | W=42 | p-value=0.15358 | r=0.467
 256 vs 1024 | W=55 | p-value=0.00592 | r=0.886
```

### S-UTIL.1 — Spectrogram PDF Catalog (utils/ai_pam_spectrogram_report.py)

#### Purpose and rationale

Visual inspection of exported spectrogram images is an essential quality-control step between Stage 4 (preprocessing) and Stage 5 (training). Anomalies such as blank images from silent segments, artifacts from boundary effects, or incorrect frequency cropping are immediately apparent on visual examination but invisible to aggregate training metrics — a model trained on a contaminated dataset may achieve plausible accuracy while learning spurious features.

In local workstation environments, spectrogram images can be browsed directly in a file manager. However, in headless computing environments — HPC clusters, SLURM-managed nodes, cloud instances — no graphical display is available. The spectrogram catalog utility addresses this gap by generating self-contained PDF documents that collect all exported spectrograms for a given class into a paginated grid layout. The resulting PDF files can be transferred to a local machine via scp or downloaded from a shared filesystem and examined with any standard PDF viewer.

#### Usage

The utility reads the spectrogram output directory from the project YAML configuration file and generates one PDF per class:

```
python utils/ai_pam_spectrogram_report.py conf/binary_cnn.yaml
```

##### Command-line options

| Option | Default | Description |
|---|---|---|
| --dpi | 150 | PDF rendering resolution. Higher values produce smaller images on the page, fitting more per sheet. |
| --classes C0 C1 | all | Restrict generation to specific classes. |
| --max-images N | all | Limit the number of images per class (useful for quick checks). |
| --max-pages N | 50 | Maximum pages per PDF file. Larger classes are split into numbered parts. |
| -v | off | Verbose mode: prints one line per image processed. |

##### Grid layout

The DPI parameter controls the physical size of each spectrogram cell on the page: a 224 × 224 pixel image occupies 224/DPI inches per side. Higher DPI values therefore produce smaller cells and denser grids. For the 224 × 224 pixel spectrograms used in this study, the approximate layout on an A4 page is:

| DPI | Cell size (approx.) | Grid (cols × rows) | Images/page | Recommended use |
|---|---|---|---|---|
| 100 | ~57 mm | 3 × 4 | ~12 | Full-size inspection of individual spectrograms |
| 150 | ~38 mm | 4 × 6 | ~24 | Balanced detail for routine quality control |
| 300 | ~19 mm | 9 × 11 | ~99 | Thumbnail overview of large classes |

Each image cell includes a truncated filename label for traceability. Page headers display the class name, page number, and total image count.

##### Output

PDF files are written to the report directory specified in the YAML configuration:

```
<report_dir>/spectrogram_catalog_C0.pdf
<report_dir>/spectrogram_catalog_C0_part2.pdf   (if exceeding max_pages)
<report_dir>/spectrogram_catalog_C1.pdf
```

##### Dependencies

This utility requires the fpdf2 library:

```
pip install fpdf2
```

The fpdf2 package is included in the project's requirements.txt and is installed automatically with a full dependency install.

#### Practical recommendations

For a typical binary detection experiment with ~8,000 C0 and ~5,600 C1 images, a DPI of 300 produces compact catalogs of approximately 60 and 40 pages respectively — easily browsable in a single PDF viewer session. At DPI 150 the same classes require roughly 230 and 160 pages, but each spectrogram is large enough to inspect frequency contours in detail.

A recommended quality-control workflow is:

1. Generate a thumbnail overview at DPI 300 with –max-images 500 to scan for systematic issues (blank images, wrong frequency range, encoding artifacts).
2. If anomalies are detected, regenerate the affected class at DPI 100 to inspect individual spectrograms at full size.
3. After confirming visual quality, proceed to Stage 5 training.

On HPC systems, the catalog generation can be included as a post-processing step in the SLURM preprocessing job:

```
#!/bin/bash
#SBATCH --job-name=pam-preproc
#SBATCH --ntasks=1 --cpus-per-task=4 --mem=16G

source ~/pam_env/bin/activate
python ai_pam_preprocessing.py conf/binary_cnn.yaml -v

# Visual QC catalog (adds ~2 min for 13,000 images at DPI 300)
python utils/ai_pam_spectrogram_report.py conf/binary_cnn.yaml --dpi 300
```

#### License

The ai-pam-pipeline software is distributed under the Apache License, Version 2.0. This permissive license allows unrestricted use, modification, and redistribution of the software — including forking in other development tree — provided that the original copyright notice and license text are retained and that any modifications are clearly marked. The full license text is included in the repository as the file LICENSE.
